# Efficiency of an allosteric protein: Multiple affinity-efficacy correlations link agonist binding to nicotinic receptor activation

**DOI:** 10.1101/2023.02.12.528209

**Authors:** Dinesh Indurthi, Anthony Auerbach

## Abstract

Receptors signal by switching between resting and active shapes under the influence of agonists. The maximum response produced by an agonist (‘efficacy’) depends on its relative binding strength (‘affinity’) to active versus resting conformations. Efficiency, the logcorrelation between these two agonist properties, is the fraction of binding energy converted into energy for the receptor’s conformational change. In adult muscle nicotinic receptors, efficiencies estimated from 76 concentration-response curves (23 agonists, 53 mutations) segregate into 5 discrete classes (%): 0.56 (17), 0.51(32), 0.45(13), 0.41(26) and 0.31(12). There is a strong linear correlation between affinity and efficacy within each class, but the multiplicity of classes precludes the appearance of any overall relationship. The efficiency class distribution indicates that there are at least 5 resting versus active binding site structural pairs. We discuss efficiency as a quantitative measure of energy coupling between agonist binding and protein conformational change that is fundamental to receptor operation.

## INTRODUCTION

The primary job of a receptor is to convert chemical energy from agonist binding into mechanical work of conformation change. Receptor theory holds that each agonist has an affinity that measures how strongly it binds to the target site, and an efficacy that measures how well the ligand activates the receptor once it is bound. Recently, a third universal agonist property, efficiency (D; eta), was introduced as the correlation between affinity and efficacy (Nayak et al., 2019). Here, we describe and interpret the distribution of efficiency values estimated from dose-response curves for different agonists and mutations of neuromuscular acetylcholine receptors (AChRs).

AChRs are five subunit, ligand-gated ion channels that have 2 neurotransmitter sites in the extracellular domain and a narrow region in the pore domain that governs the passive flow of cations across the membrane (Gharpure et al., 2020; Unwin, 1995; Zarkadas et al., 2022). In activation, and to a first approximation, AChRs alternate between a resting closed-channel (C) and an active open-channel (O) conformation, stable shapes in which the neurotransmitter sites bind agonists either weakly (low affinity, LA) or strongly (high affinity, HA). The neurotransmitter sites and the pore constriction are separated by ∼60 Å, but both regions change structure and function conjointly within a global ‘gating’ isomerization, C_LA_⇄O_HA_.

In AChRs and many other receptors, without any bound agonists the probability of the active-O conformation (P_O_) is small and baseline activity is negligible (Jackson, 1986; Nayak et al., 2012; Purohit & Auerbach, 2009). However, when both agonist sites are occupied by the neurotransmitter acetylcholine (ACh) the extra favorable binding free energy to O (compared to C) increase P_O_ and membrane current substantially (Figure 1). Indeed, the low- and highconcentration asymptotes of the concentration(dose)-response curve (CRC) for ACh and wildtype AChRs are approximately 0 and 1.

**Figure 1.**
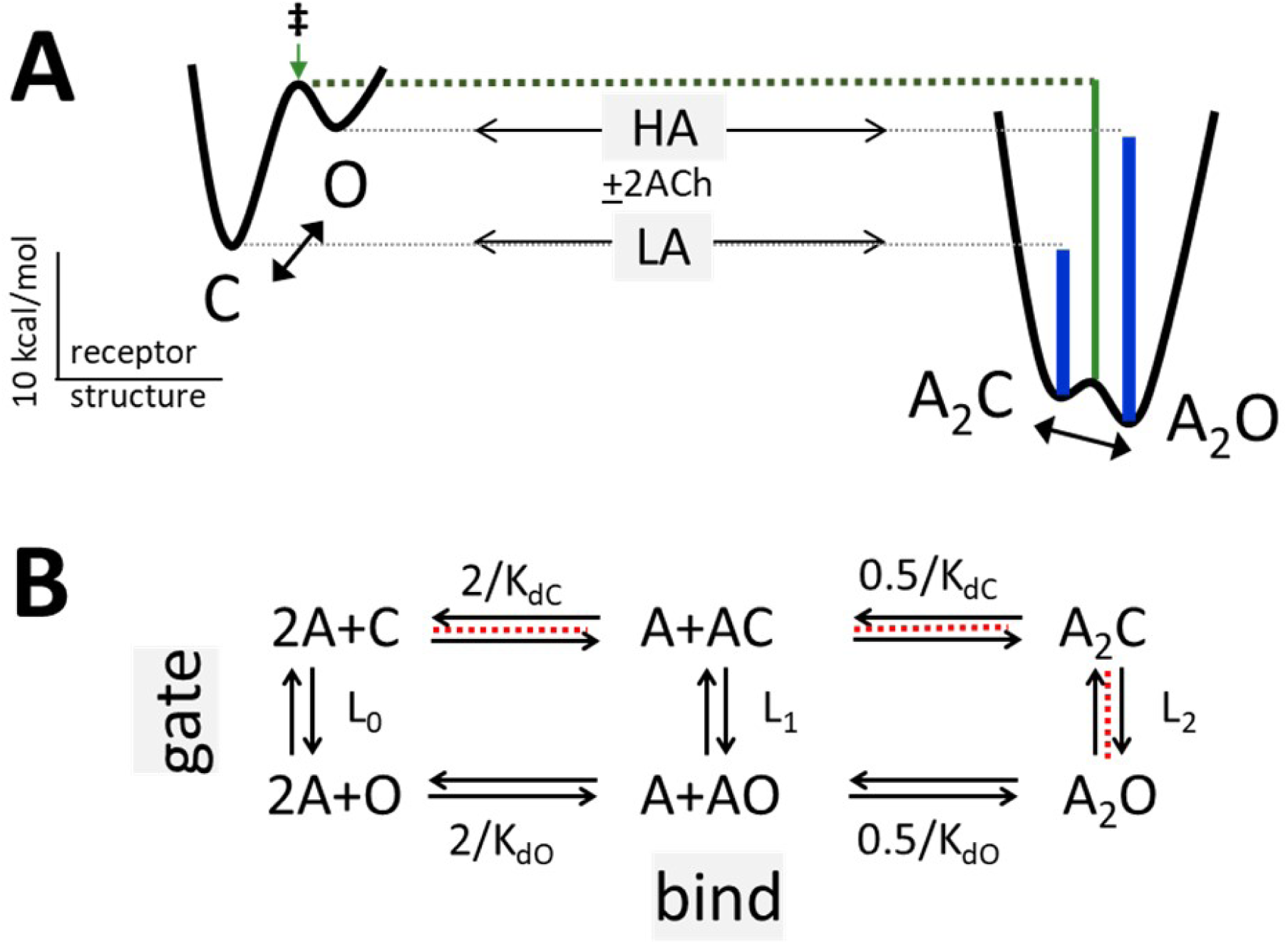
Bind and gate. A. Potential energy surfaces (landscapes) for activation without (left) and with (right) bound agonists (A). C, closed channel and low affinity (LA); O, open channel and high affinity (HA). Agonists increase the probability of O (P_O_) because they bind more strongly to their target sites in O versus C (vertical blue lines). green, agonists bind with HA to the gating transition state (‡). calibration: ACh, adult-type AChRs, -100 mV, 23 °C. B. Corresponding reaction scheme (2 equivalent and independent neurotransmitter sites). horizontal (bind): K_dC_ and K_dO_, LA and HA equilibrium dissociation constants to C and O; vertical (gate): L_n_, equilibrium constant with n bound agonists. L_0_ is agonist independent; red, main pathway for concentration-response curves (Eq. 4, Methods).

Our approach to understanding how agonists activate AChRs is to estimate the free energy changes associated with each step in the process. Briefly, we use electrophysiology to compile a CRC for each agonist-receptor combination, and from its asymptotes and midpoint estimate both equilibrium dissociation constants (K_dC_ and K_dO_) and, hence, the corresponding weak and strong binding free energies (DG_LA_ and DG_HA_). As shown below, agonist efficiency is calculated from these energies.

Previously, it was observed that in adult-type AChRs the weak/strong ratio of binding energies, DG_LA_/DG_HA_, is approximately the same for a group of agonists despite wide variations in individual agonist affinity and efficacy (Jadey & Auerbach, 2012). Within this group, whose members include full agonists (like ACh), partial agonists (like nicotine) and very weak agonists (like choline), the binding energy ratio is ∼0.50. That is, for all members of this group binding to O is ∼2-fold stronger than to C, regardless of affinity or efficacy.

Subsequently, two additional agonist groups were identified having binding energy ratios of 0.58 (for example, epibatidine) and 0.46 (for example, tetramethylammonium) (Indurthi & Auerbach, 2021; Nayak et al., 2019). For these, agonist binding free energy increases by 1.7- and 2.2-fold in the channel-opening process. Efficiency depends only on the binding energy ratio (Jadey & Auerbach, 2012), with the abovementioned 3 agonist groups corresponding to D values of 0.54, 0.50, 0.42.

The observation of a common DG_LA_/DG_HA_ ratio for even one group of ligands is interesting because it indicates that the only two agonist-dependent events in the activation process - weak binding to C and the switch to strong binding that happens within gating - are not independent. Rather, a shared binding energy ratio for a group of ligands implies that the structure (energy) changes that underpin affinity (part of binding) and efficacy (part of gating) are linked in a linear free energy relationship (LFER) (Auerbach, 2016; Jadey & Auerbach, 2012; Purohit et al., 2014). The relationship between affinity and efficacy has been considered previously (Colquhoun, 1998; Kenakin, 2002), although not with regard to weak versus strong agonist binding energies and this LFER.

Empirically, efficiency is simply the correlation between affinity and efficacy. Further, because of the LFER, efficiency is the fraction of binding energy that is converted into energy for the receptor’s gating conformational change. The parameter D, calculated from a single CRC, calibrates the strength of coupling between receptor input and output.

We have measured D for various AChR agonists and mutations. We report 16 new values that, combined with previous measurements, describe a spectrum of 5 D classes that conceals the underlying correlations between affinity and efficacy that are apparent within each class. Multiple binding energy ratios (efficiency classes) indicates there are multiple resting versus active binding site structural pairs, and, possibly, multiple subtypes of gating isomerization. Importantly, the existence of efficiency classes shows that binding and gating are energy-linked stages of a unified allosteric transition.

## RESULTS

<u>Theory.</u> The standard conception of receptor activation incorporates two apparently independent steps, bind and gate (Figure 1B). In bind the agonist (A) diffuses to the resting target site and forms a LA complex (A+C⇄AC_LA_), and in gate agonist binding strength increases as part of the global isomerization to the open channel, HA shape (AC_LA_⇄AO_HA_). The global gating isomerization is neither instantaneous nor synchronous but rather involves passage through short-lived (<Ds) intermediate conformations (Figure 2 – figure supplement 1). In AChRs, the affinity increase occurs near the start of the channel-opening isomerization (Grosman et al., 2000; Purohit et al., 2013) so the increased agonist binding free energy is almost fully realized at the gating transition state (‡) as well as at the final O state. Hence, agonists increase P_O_ almost exclusively by increasing the opening rate constant.

Affinity and efficacy are universal agonist attributes. Regarding affinity, receptors are allosteric proteins that have 2 ligand binding strengths, DG_LA_ and DG_HA._ In AChRs, both of these have been measured directly and independently (Nayak & Auerbach, 2017). Efficacy can be defined in several ways. First, it is the high-concentration asymptote of an unnormalized CRC (P_O_^max^). Considering just bind and gate, this limit depends only on the fully-liganded gating equilibrium constant, L_2_ (Figure 1B; Eq. 4a). A second definition derives from considering the full cycle of receptor activation. Experiments show that in adult-type AChRs the 2 neurotransmitter binding sites are approximately equivalent and independent, and that there is no significant input of external energy (Nayak & Auerbach, 2017). Hence,

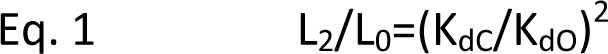

where the gating equilibrium constant subscripts refer to the number of bound agonists and the equilibrium dissociation constant ratio is the ‘coupling’ constant (c). L_0_ is agonistindependent, so differences in L_2_ among agonists are determined only by c. A third definition is the logarithm of the coupling constant, DG_HA_-DG_LA_. P_O_, L_2_, c and this binding energy difference are all measures of relative agonist efficacy.

We now define agonist efficiency as the ratio of efficacy (binding energy difference) and affinity (strong binding energy to O):

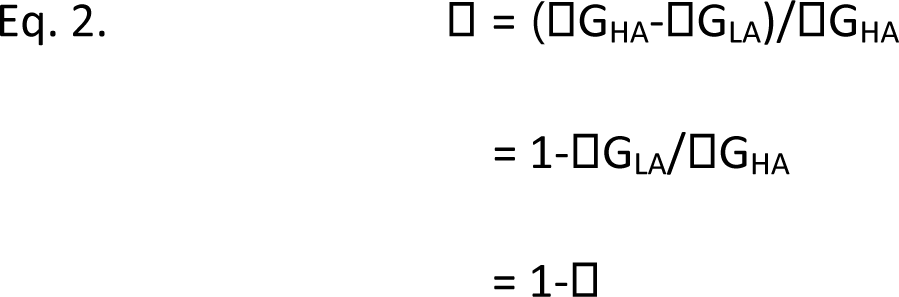

where D is the ratio DG_LA_/DG_HA_. This empirical definition does not imply that there is one efficiency value for all agonists, only that an D-value can be calculated for any agonist from K_dC_ and K_dO_. We estimated these equilibrium quantities from CRCs, but any experimental approach that measures K_dC_ and K_dO_ would suffice.

The above definition of efficiency is empirical, but there also is a physical interpretation. In AChRs, both bind and gate are composite reactions (Figure 2 – figure supplement 1) (Auerbach, 2005; Gupta et al., 2017; Jadey & Auerbach, 2012). Binding involves both agonist diffusion to, and a local rearrangement of, the neurotransmitter site (‘catch’). Gating involves both a second binding site rearrangement (‘hold’) and additional conformational changes in other protein domains (Figure 2A). As far as agonist action is concerned, only the catch and hold microscopic transitions within bind and gate are relevant. The agonists we examined all have a similar diffusion constants and post-hold domain rearrangements that are mostly agonist independent (Gupta et al., 2017). Therefore, with regard to the activation sequence that determines the CRC, namely A+C⇄AC⇄A_2_C⇄A_2_O (red, Figure 1B), differences between agonists are caused mainly by differences in the free energy changes in catch and hold (Figure 2B).

**Figure 2.**
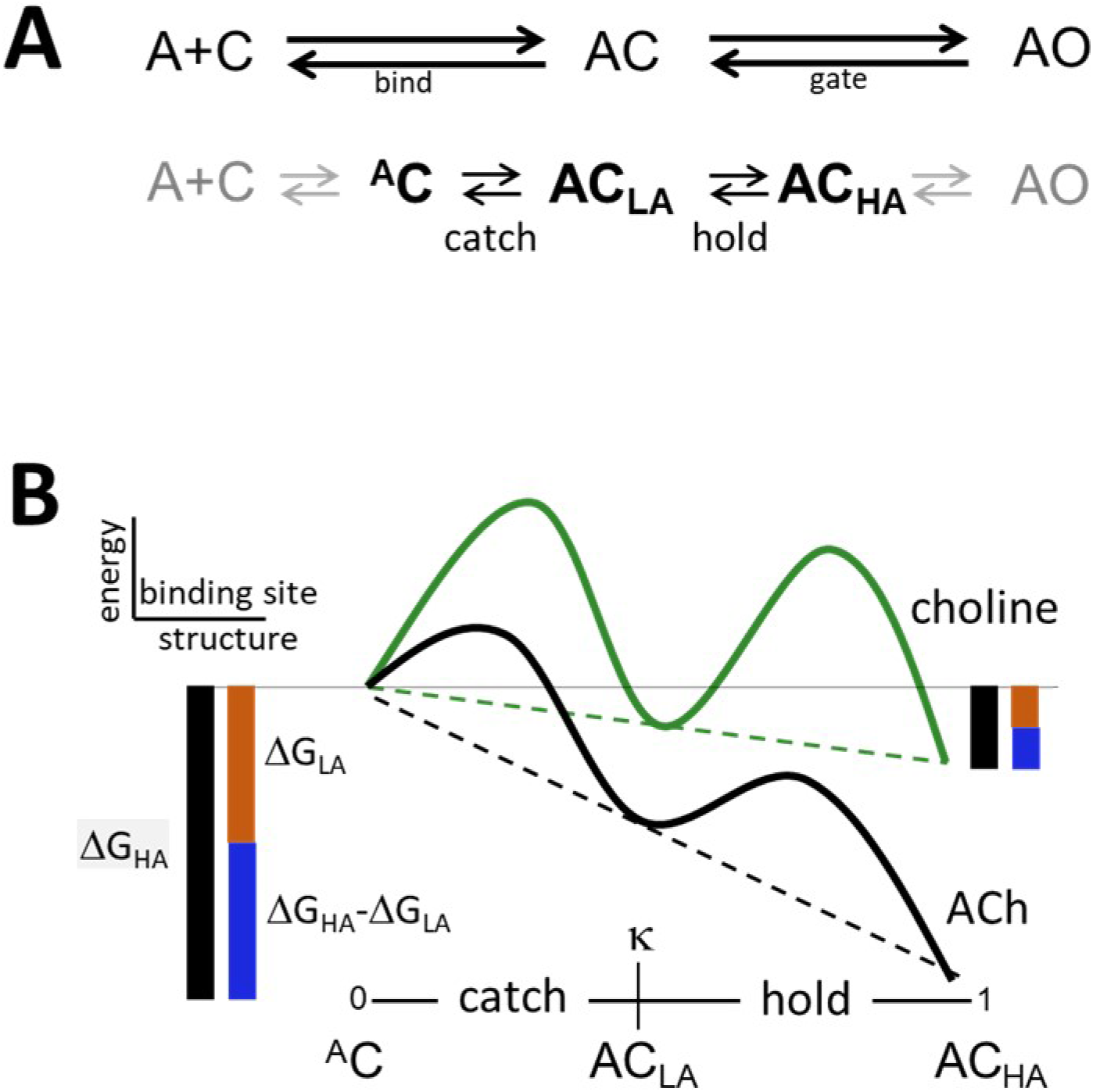
Catch and hold. A. Catch and hold are intermediate steps in bind and gate; gray, agonist-independent steps (see Figure 2 – figure supplement 1). B. Catch-and-hold free energy landscapes. The two binding site rearrangements (^A^C⇄AC_LA_ and AC_LA_⇄AC_HA_) are linked in a linear free energy relationship (LFER; dashed lines); green, partial agonist; black, full agonist. There are 3 agonist-protein complexes: ^A^C (post-diffusion), AC_LA_ (post-catch) and AC_HA_ (posthold). Agonists tilt the LFER to a total extent DG_HA_ (black side bars). DD fraction of the total tilt expended in catch (affinity; brown bar); DD the remainder expended in hold (efficacy; blue bar). Affinity and efficacy of partial versus full agonists differ, but their efficiency is the same because it only depends only on the position of AC_LA_ in the reaction co-ordinate (see last paragraph, Methods).

After removing the free energy associated with agonist diffusion (see next paragraph), the residual free energy changes that can be calculated from the logarithms of the equilibrium dissociation constants are DG_LA_ (catch) and DG_HA_-DG_LA_ (hold) (Figure 2B). Their sum, DG_HA_, is the total free energy change experienced by the agonist in the activation process. Accordingly, the above empirical definitions of D and D (Eq. 2) can be interpreted as the fractions of binding energy change associated with catch and hold. Multiplying by 100% gives these as percentages. D is our primary focus because it pertains to the binding site rearrangement that jumpstarts the cascade of domain rearrangements that constitute gating.

The above physical interpretations of D and D could still be ambiguous because the component binding free energies include a chemical potential that has an entropy term indexed to a standard ligand concentration (Dill. K., 2010). We removed this potentially confounding element by first normalizing each of the component equilibrium dissociation constants to those for a standard ligand (Methods). As shown below, D values calculated without normalization are <5% greater than with normalization, indicating that the free energy changes associated with the chemical potential are small compared to those associated with the formation of the ligand-protein complex. Hence, the entropy offsets associated with ligand binding *per se* are minimal, and D is a true ratio of free energies. D represents the fraction of the total ligandprotein binding free energy associated with catch, and D is the remainder associated with hold. Note that D gives the fractional energy change at the middle state in 2-step LFER (at an energy well; DG_LA_/DG_HA_), whereas the parameter D (or D) gives the fractional energy change at the transition state in a 1-step LFER (at an energy barrier; DG^‡^/DG^0^) (see last paragraph, Methods).

EC_50max_ P_o_ are observed values estimated from each CRC (source data in Figures. 3 and Figure 3 – figure supplement 1). K_dC_ and K_dO_ are calculated values that have been corrected for background mutations that only change L_0_ (Eq. 4). c, coupling constant (K_dC_/K_dO_); D, efficiency (1-logK_dC_/logK_dO_) (Eq. 2); n, number of CRCs. Membrane potential, +70mV. Agonist (Figure 4) superscripts are the backgrounds: ^a^εS450W, ^b^εL269F, ^c^εE181W. sem, standard error of mean.

<u>Agonists.</u> We measured D and D in AChRs having wild-type (wt) binding sites for 5 previously unstudied agonists (Figure 3, Figure 3 – figure supplement 1, Table 1). Single-channel currents were recorded at different agonist concentrations and P_O_ values calculated from shut and open interval durations were compiled into a CRC. L_0_ was known *a priori* (Nayak et al., 2012). As shown elsewhere, CRCs complied from whole-cell currents serve equally well (Indurthi & Auerbach, 2021).

**Figure 3.**
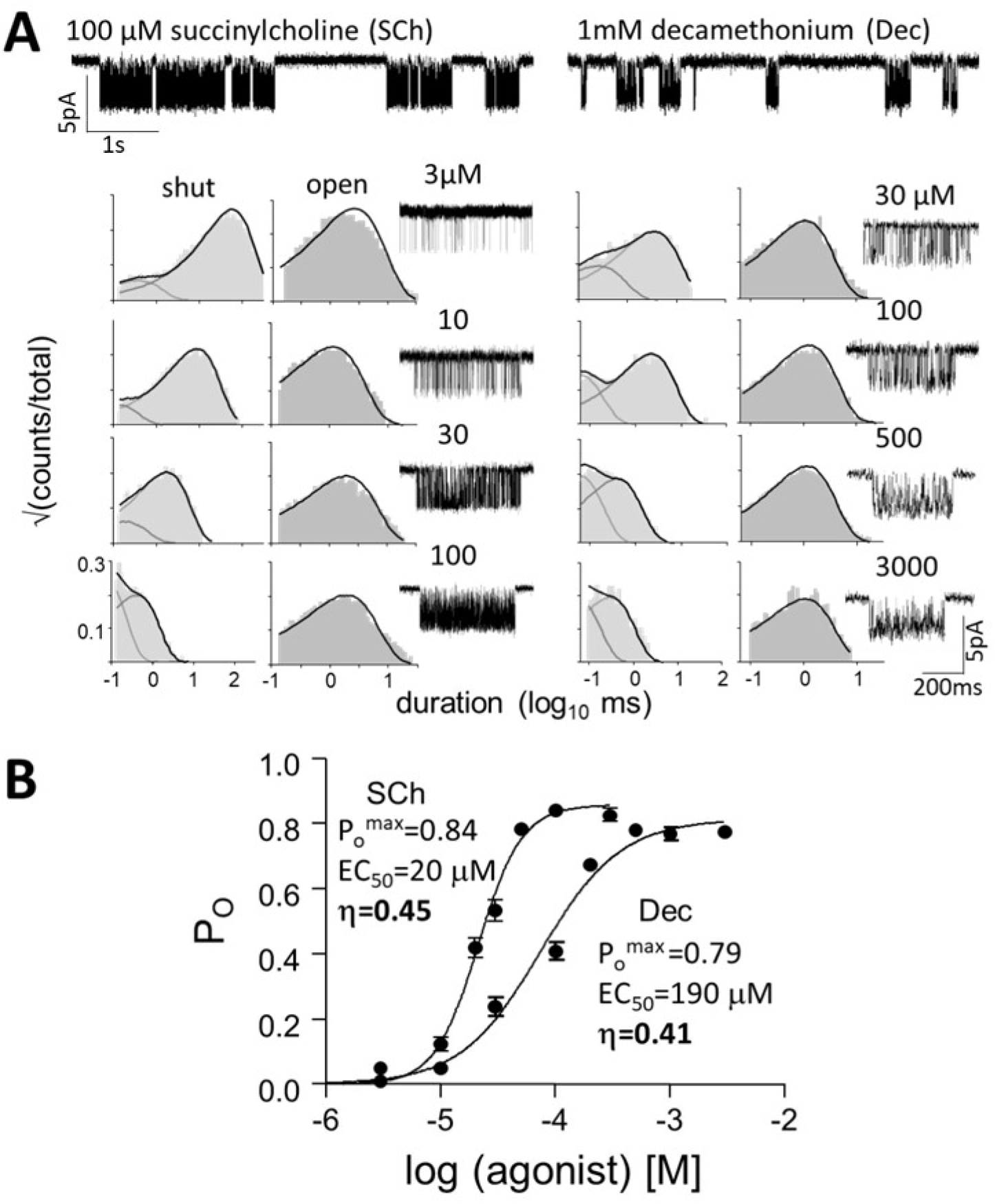
Measuring agonist efficiency. A. Top, single-channel current traces at low timeresolution (O is down). Clusters of openings are bind-gate; silent periods between clusters are desensitized. Bottom, example intra-cluster interval duration histograms and clusters. Cluster P_O_ was calculated from the shut- and open-interval time constants at each [agonist]. B. CRCs. P_O_ values were fitted to estimate P_O_^max^ and EC_50_ from which K_dC_ and K_dO_ were calculated (Eq. 4, Methods) to yield agonist efficiency (D) (Table 1). The profile for SCh is relatively left-shifted because this agonist is ∼10% more efficient than Dec. Symbols are mean+sem. See Figure 3 – figure supplement 1 for similar results regarding other agonists used in this study.

Figure 3 shows CRCs for succinylcholine (SCh) and decamethonium (Dec). Although P_O_ is similar for both agonists, the EC_50_ (potency) of SCh is substantially lower than that of Dec. For each agonist K_dC_ and K_dO_ were estimated from the CRC parameters (Eq. 4), and DG_LA_ and DG_HA_ were calculated as being proportional to logK_dC_ and logK_dO_, to yield D by using Eq. 2 (Table 1). The results were D_SCh_=0.45 and D_Dec_=0.41. SCh is ∼10% more efficient than Dec, which is the root cause of the abovementioned mismatch between P_O_ and potency. Given L_0_, agonist efficiency can be calculated from a single dose-response curve. Every agonist of every receptor has an affinity, efficacy and efficiency. Errors associated with D estimated from P_O_^max^ and EC_50_ are small and are discussed elsewhere (Indurthi & Auerbach, 2021).

**Table 1.**
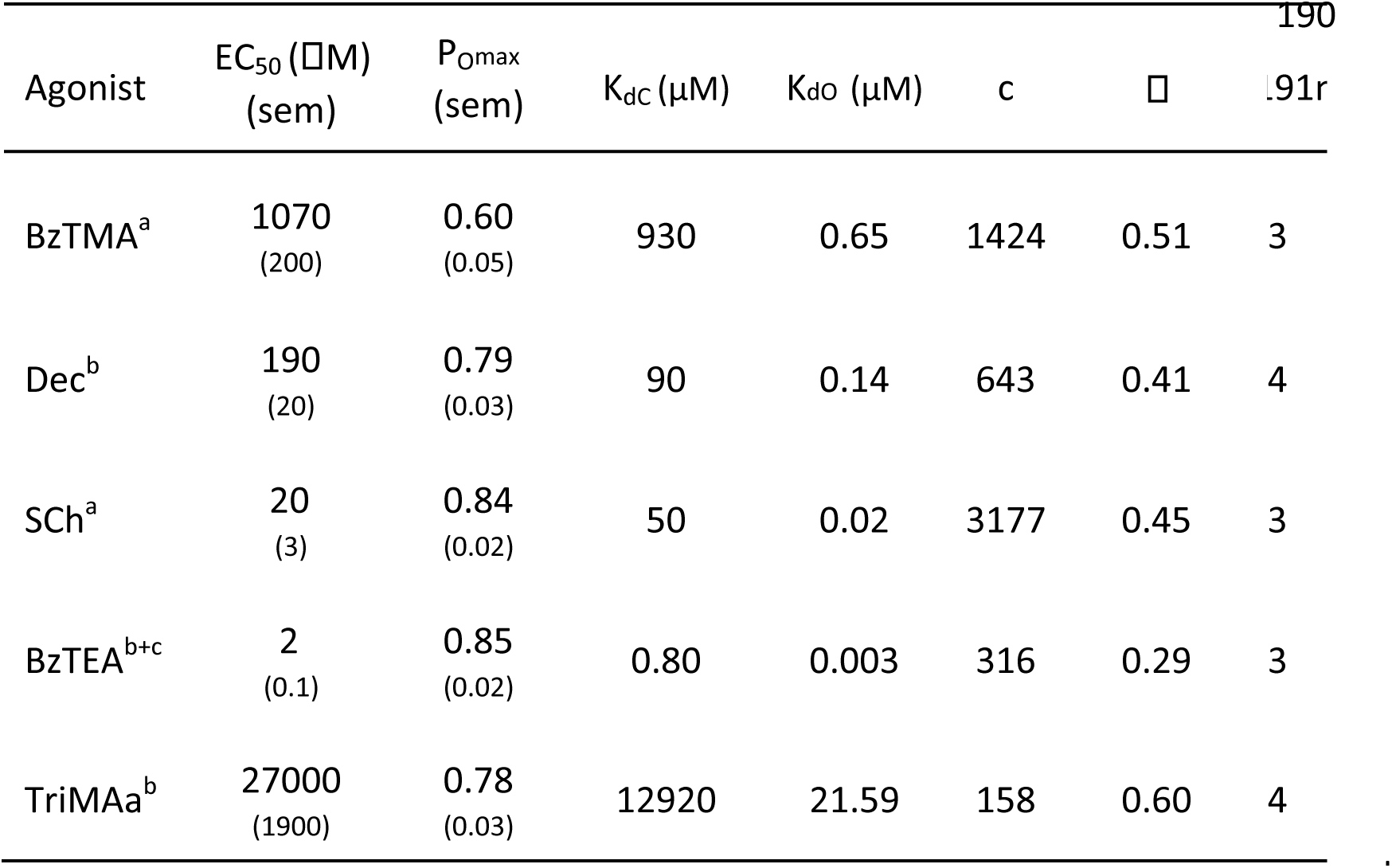
Agonist efficiencies

So far, D has been estimated for 23 agonists at wt D−DDD (adult) AChR binding sites (Figure 4, Table 1 and Table 1 – table supplement 1). Figure 5A (main) shows a plot of weak versus strong binding energies (slope D) for these ligands, absent the chemical potential. The points segregate into 5 different straight lines (using x-means clustering, see methods), each of which represents an efficiency class. Figure 5A (inset) shows that D values estimated without excluding the chemical potential are only slightly greater.

**Figure 4.**
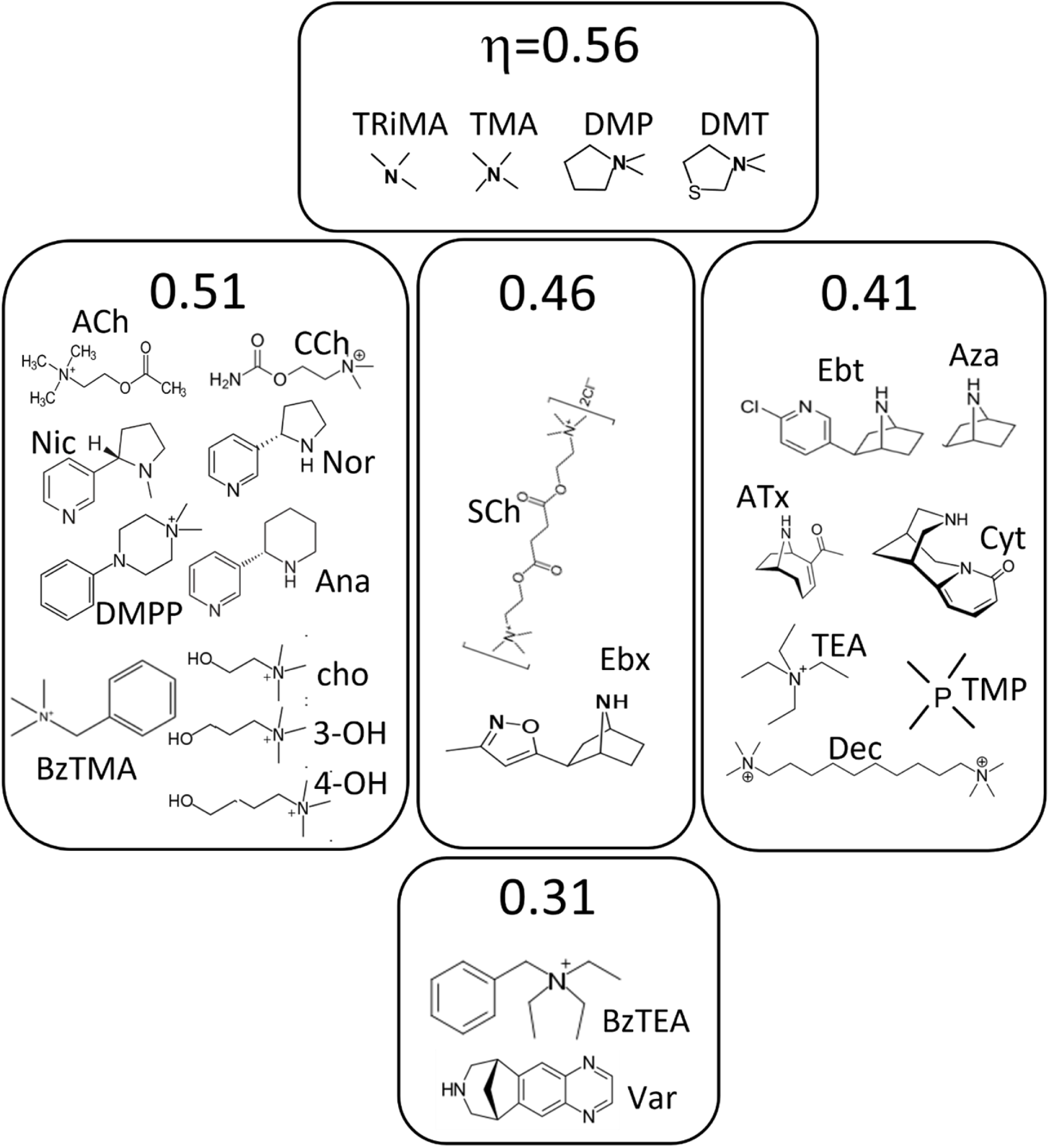
Agonists. Ligands are grouped by efficiency class (Figure 5B). See Methods for abbreviations.

**Figure 5.**
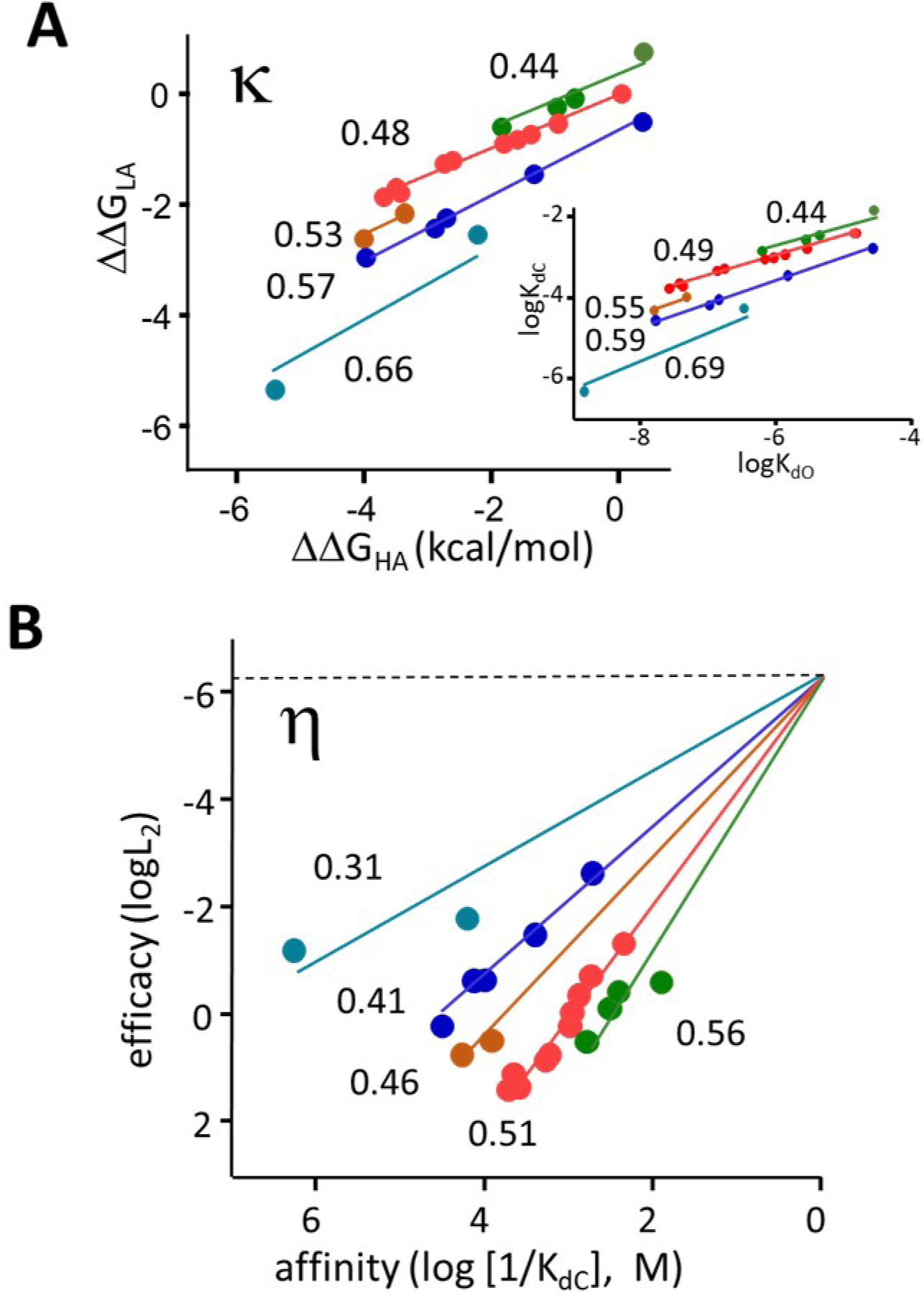
Agonist efficiencies. A. D-plot. Each symbol represents a different agonist, weak binding energy to C (proportional to logK_dC_) versus strong binding energy to O (proportional to logK_dO_). Main panel, K_d_s were normalized to those for choline to remove the chemical potential. Inset, unnormalized values. Slopes without normalization are <5% greater. There are 5 slopes (D values) (see Figure 7A inset)D A D-plot for a 2 step LFER (Figure 2B) is analogous to a D-plot for a 1-step LFER. B. D-plot (log affinity versus log efficacy). Log axes are binding versus gating equilibrium constants (Figure 1B), proportional to DG_LA_ versus (DG_HA_-DG_LA_), or energy changes in catch versus hold (Figure 2B). The points were fitted by Eq. 3 using L_0_=5.2×10^-7^. D was calculated from each slope. There are 5 agonist efficiency classes (Figure 4).

Although D can be calculated directly as 1-D (Eq. 2), we used an alternative approach. Combining Eqs. 1 and 2 (Nayak et al., 2019),

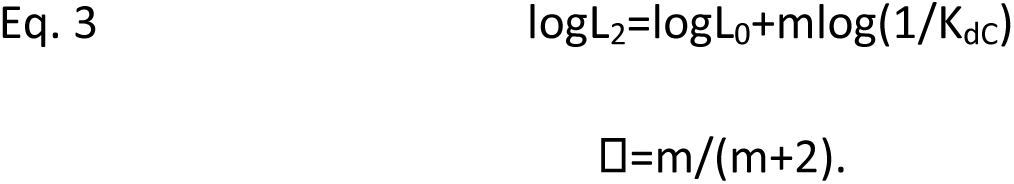

A log-log plot of L_2_ (y-axis) versus 1/K_dC_ (x-axis), or efficacy versus affinity, is an ‘efficiency’ plot. A fit by a straight line gives an estimate of D from the slope and L_0_ from the y-intercept. Eq. 3 converts readily measured composite parameters of a CRC (P_O_ and EC_50_) into fundamental constants of agonists and receptors (D and L_0_) *via* equilibrium constants (K_dC_ and L_2_) (Eq. 4). Here, the main advantage of using the D-plot is that, because L_0_ is the same for all agonists and has been measured previously, the y-intercept can be added as a fixed point to all lines to improve the accuracy of the slope estimates. For receptors in which L_0_ has not been measured, the efficiency plot offers a convenient way to do so from CRCs of agonists that have the same D.

Figure 5B shows D-plots for all 23 agonists that have been examined so far. From the slopes, we estimate that the mean values of the 5 D classes to be 0.31, 0.41, 0.46, 0.51 and 0.57. In Figure 4 the agonists are grouped by D. The neurotransmitter ACh, its breakdown product choline, the partial agonist carbamylcholine (CCh) and nicotine all belong to the 0.51 class. The second-most common class, 0.41, includes ligands that have a bridge nitrogen (for instance, Ebt) and others that do not (for instance, Dec and TMP). The smallest ligands (for instance, TriMA and TMA) were the most efficient, and the largest (rigid) ligands (varenicline, BzTEA) were the least efficient.

The efficiency plot shows the correlation between affinity and efficacy on a log-log scale. Figure 5B indeed shows that agonists with the same affinity can have different efficacies and *vice versa* (Pearson’s correlation confirms the correlation strength between affinity and efficacy within D classes containing more than two data points, see methods), and that in AChRs there is no apparent global correlation between these properties (data not shown). However, the lines in Figure 5B show clearly that agonists within each efficiency class do have the same correlation. In AChRs, a single correlation between affinity and efficacy is precluded by an abundance of efficiency classes.

<u>Mutations.</u> Agonist efficiency was measured previously in adult-type AChRs having one of 42 binding site mutations (Table 2 – table supplement 1). To this we add 11 values (Table 2), for 5 agonists and the mutations DD200A or DK145A, plus Ebt/DG153S (Figures 6, Figure 6 – Source Data 1, Figure 6 – Source Data 2). DD200 and DK145, along with DY190, have been suggested to work together to initiate the channel-opening conformational change (Mukhtasimova et al., 2005). The mutation DY190A reduces ACh efficiency from 0.50 to 0.35, but DY190F is without effect (Bruhova & Auerbach, 2017). However, DY190F does cause a substantial loss in DG_LA_ for ACh to an extent that depends on the DK145 side chain (Bruhova & Auerbach, 2017). The agonists we tested with DD200A or DK145A were from 4 different D classes (wt class value): TMA (0.54), CCh (0.52), ACh (0.50), Ebx (0.46) and Ebt (0.41).

**Table 2:** Mutation efficiencies.

| mutation | agonist | EC<sub>50</sub><br>(μM)<br>(sem) | P<sub>Omax</sub><br>(sem) | K<sub>dc</sub><br>(μM) | K<sub>do</sub><br>(μM) | c | η | n |
| --- | --- | --- | --- | --- | --- | --- | --- | --- |
| D200A | ACh<sup>a</sup> | 110<br>(10) | 0.80<br>(0.02) | 52 | 0.15 | 338 | 0.37 | 3 |
|  | CCh<sup>a</sup> | 320<br>(70) | 0.45<br>(0.02) | 138 | 0.90 | 153 | 0.36 | 4 |
|  | TMA<sup>a</sup> | 13350<br>(2970) | 0.37<br>(0.03) | 5683 | 43.82 | 130 | 0.48 | 5 |
|  | Ebt<sup>a+b</sup> | 110<br>(20) | 0.50<br>(0.03) | 47 | 0.68 | 69 | 0.30 | 3 |
|  | Ebx<sup>a+b</sup> | 060<br>(20) | 0.49<br>(0.03) | 26 | 0.39 | 67 | 0.29 | 3 |
| K145A | ACh<sup>a</sup> | 110<br>(10) | 0.95<br>(0.06) | 83 | 0.05 | 1672 | 0.44 | 3 |
|  | CCh<sup>a</sup> | 380<br>(30) | 0.53<br>(0.01) | 165 | 0.40 | 411 | 0.41 | 4 |
|  | TMA<sup>a</sup> | 2200<br>(590) | 0.19<br>(0.01) | 922 | 5.01 | 184 | 0.43 | 3 |
|  | Ebt<sup>a</sup> | 40<br>(4) | 0.81<br>(0.04) | 20 | 0.02 | 792 | 0.38 | 3 |
|  | Ebx<sup>a</sup> | 40<br>(3) | 0.83<br>(0.01) | 20 | 0.02 | 847 | 0.38 | 3 |
| G153S | Ebt<sup>a</sup> | 2<br>(0.5) | 0.65<br>(0.01) | 2 | 0.007 | 313 | 0.31 | 4 |

**Figure 6.**
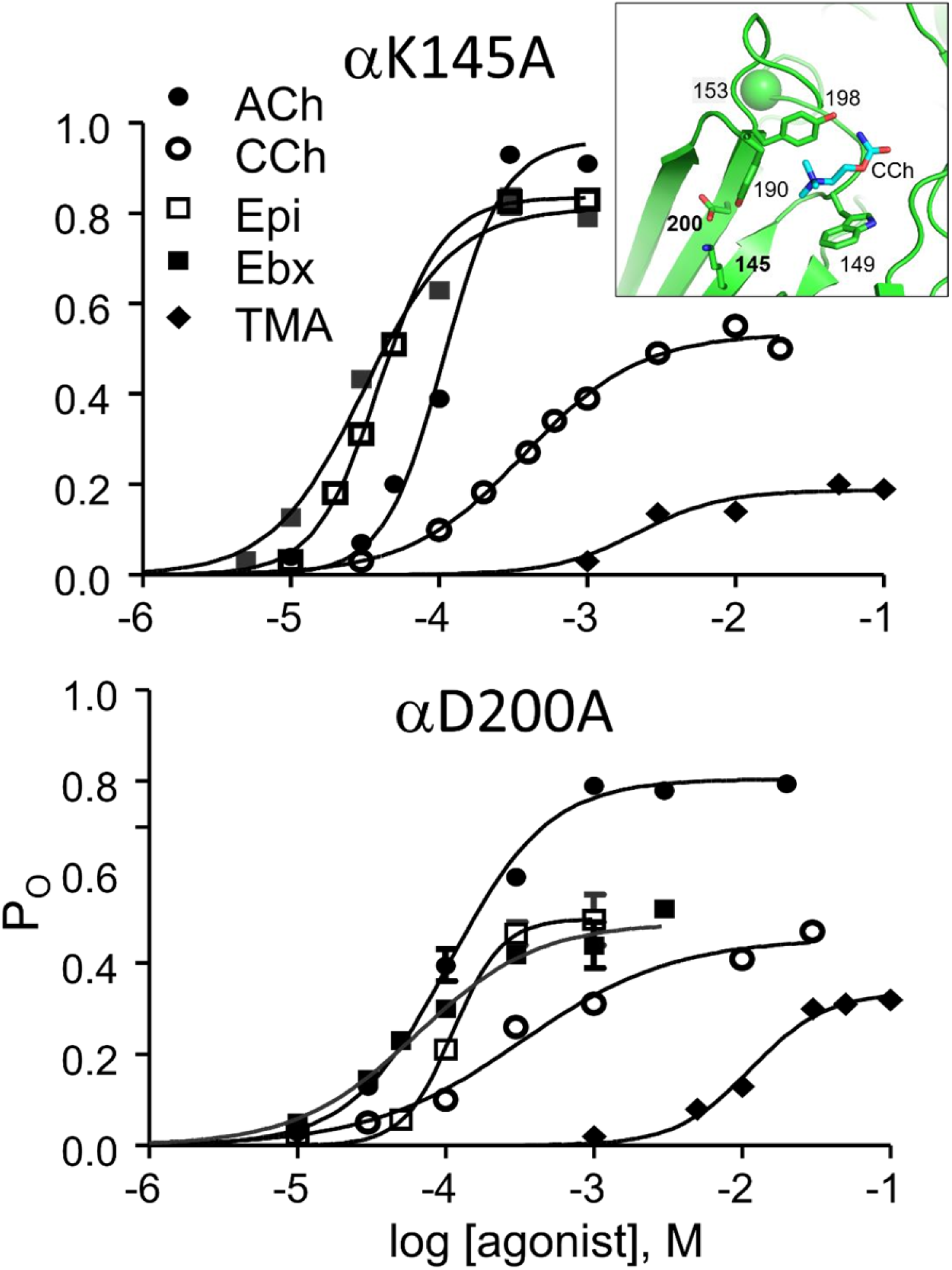
CRCs for DK145A and DD200A. Inset, D subunit of an AChR neurotransmitter binding site (7QL6.pdb; (Zarkadas et al., 2022)). Efficiencies calculated from the fitted CRC parameters are in Table 2. For intra-cluster interval histograms see Figure 6 – Source Data 1 and Figure 6 – Source Data 2.

For mutation location see Figure 6 inset; wt agonist efficiencies given in Figure 4. K_dC_ and K_dO_ were calculated from CRC parameters (Figure 6) by using Eq. 4. c, coupling constant (K_dC_/K_dO_); D, efficiency (1-logK_dC_/logK_dO_); n, number of CRCs. L_0_ was corrected for background mutations (Methods): ^a^εS450W, ^b^εL269F, ^c^εE181W. See Figure 6 – Source Data 1 and Figure 6 – Source Data 2 for source data.

At DD200, an A substitution reduced the average efficiency of 4 of the 5 agonists to 0.33D0.04 (mean±sd), or to about the same level as ACh with DY190A. The exception was TMA for which efficiency remained high at 0.48. D for all 5 agonists was 0.41D0.03 following an A substitution at DK145. Interestingly, this mutation reduced D for ACh, CCh and TMA but did not significantly alter the efficiencies of Ebx and Ebt.

Table 2 – table supplement 1 shows K_dC_ and K_dO_ values measured previously for 4 different agonists in AChRs having a substitution at DG153 (Jadey et al., 2013), including a Ser that causes a congenital myasethetic syndrome (Engel et al., 1982). After converting to DG_LA_ and DG_HA_, these results indicate that on average the mutations reduce D for choline, DMP, TMA and nicotine to 0.41+0.03 (mean+sd), or by ∼25%. In addition, we now report that DG153S reduces the efficiency of Ebt from 0.42 to 0.31, or also by ∼25%.

With mutations, an D-plot is no more useful than a D-plot because substitutions can change L_0_ (see Figure 7 – figure supplement 1) and, hence, the y-intercept of each line. Figure 7A shows a D-plot for all mutations that have been measured so far. Efficiency estimates for mutations have more scatter than with agonists because of the requirement that L_0_ be measured independently for each side chain substitution. Nonetheless, D values calculated from D values after a mutation segregate into the same 5 classes (Figure 7A, inset) that were apparent with agonists (Figure 5B), with significant correlation strength between low and high binding affinities within each class, see methods. Considering both agonists and mutations, the efficiency value (relative prevalence) of each class is 0.56 (17%), 0.51(31%), 0.45(13%), 0.41(26%) and 0.31(12%) (Figure 7B). As was the case with just agonists, the 0.51 and 0.41 efficiency classes predominate. Figure 7C shows the distribution of agonist plus mutation D values as a spectrum or ‘bar code’ in which line thickness represents relative prevalence.

## DISCUSSION

The main results are as follows. 1) Empirically, efficiency is the log-log correlation between affinity and efficacy. It is a universal parameter that applies to every agonist of every receptor. 2) Physically, efficiency is the fraction of agonist binding energy associated with the local rearrangement of the binding site that triggers gating. Efficiency depends only on the agonist’s weak/strong binding free energy ratio. 3) In AChRs, agonists with radically different affinities and efficacies can have the same efficiency because the ligand’s free energy changes in binding and gating are linked in a LFER (catch and hold). 4) There are 5 agonist efficiency classes in AChRs that together disallow a global affinity-efficacy correlation. 5) Mutations of binding site residues segregate into the same 5 efficiency classes as do agonists. The observation of even one efficiency class indicates that the catch-hold rearrangements within bind-gate comprise a single LFER (Figure 2). Regardless of the extent to which a ligand may ‘tilt’ the binding energy landscape to alter affinity and efficacy, the fraction of the total binding energy that is consumed in the second, gating step (hold) remains constant. Only the left-right position of the intermediate state AC_LA_ in the overall reaction co-ordinate determines efficiency. Perturbations may interfere with one or more post-hold domain rearrangements in gating, to decouple agonist binding from channel opening (Cymes & Grosman, 2021). Regardless, an efficiency calculated from K_dC_ and K_dO_ remains a measure of the fraction of binding energy applied to the hold rearrangement.

The LFER offers an insight into receptor operation. Catch and hold, and therefore bind and gate, are not independent processes but rather are entangled in a single ‘sweep’ of conformational change. In AChRs, bind-gate activation is a reversible, energy-coherent conformational cascade that links an agonist landing at the binding site to water (an ion) traversing the pore constriction. The parameters that define the free energy changes experienced by the agonist within this cascade are D and DD which are, respectively, the fraction used to form the LA complex (within bind) and the fraction associated with the switch from LA to HA (within gate). Efficiency illuminates the core of receptor operation, namely energy coupling between agonist binding and the protein conformational change that eventually turns on function.

In adult muscle AChRs there are 5 distinct D classes but there could be more. For instance, the scatter in the lowest and highest classes (Figure 7) suggests that these could be amalgams. Nonetheless, the ability to corral 76 different agonist-receptor combinations into a handful of efficiency classes represents a significant reduction in complexity. If nothing else, D is a useful way to classify agonists and binding sites.

**Figure 7.**
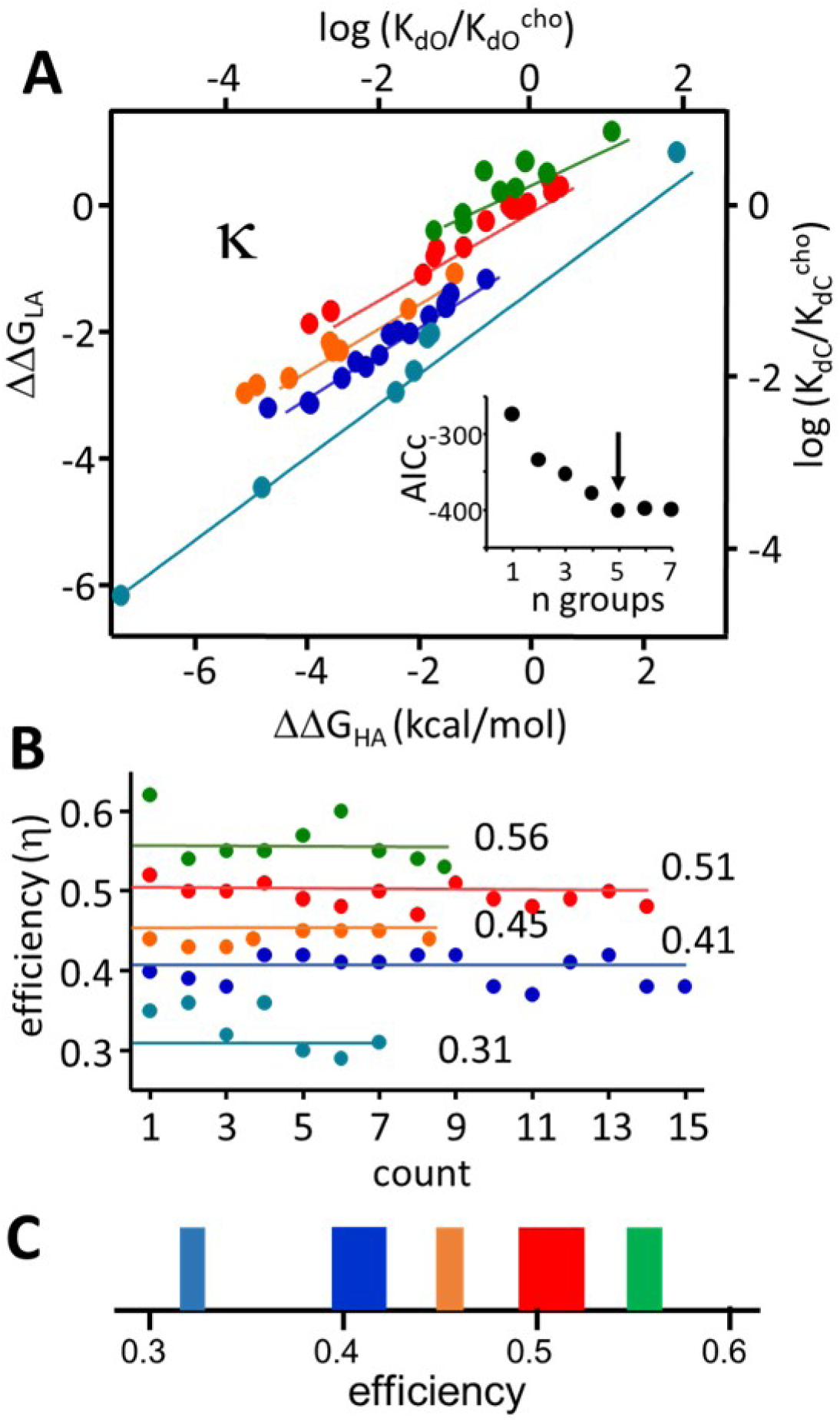
Mutation efficiencies. A. D-plot, with binding energies normalized to remove the chemical potential (see Figure 5A). Inset: corrected Akaki information criterion (AICc) indicates that the most likely number of classes is 5 (arrow), see methods for . B. Mutation efficiencies (calculated as 1-D) are approximately the same as agonist efficiencies (Figure 5B). 0.51 and 0.41 classes predominate. C. Spectrum of efficiencies, adult D−DDD neurotransmitter binding sites. The width of each line is proportional to its prevalence (agonists and mutations). D∼0.51 (Dmitrij Ljaschenko) for ACh, CCh and choline but ∼0.56 (green) for TMA. Not shown, at the fetal D−D site D∼0.56 for all (Nayak et al., 2019). See Figure 7 – figure supplement 1 and methods for L_0_ values in this study.

The distribution of D classes is the same for both agonists and mutations. An agonist at a binding site behaves much like an ordinary side chain that is exceptional only insofar that it is not (necessarily) an amino acid and not linked covalently, allowing it to be both hypermobile in the pocket and free to come and go and serve as a signal. In addition, and by definition, a agonist that is bound is more stable in O versus C. This is the opposite of most AChR side chain substitutions that typically increase L_0_, indicating that wt side chains are in general relatively more stable in C versus O. However, experiments show that at the binding sites most mutations have little effect on unliganded activity (Purohit et al., 2014).

Energy differences reflect structural ones and while these are still not known, some inferences can be made. The existence of 5 discrete D classes indicates that there are at least this many different AC_LA_-AC_HA_ binding site structural pairs, for example 1 LA+5 HA (or v*ice versa*), 2 LA+3 HA, and so on. The consistent and discrete distribution of D classes suggests that the AC_LA_ and AC_HA_ structural pairs, too, are consistent and discrete rather than continuous. The observation of 5 D classes suggest an allosteric mechanism that combines elements of conformational selection and induced fit insofar as the system selects between 5 possible discrete binding site conformation pairs depending on agonist structure (Hammes et al., 2009).

The binding energy change (hold) occurs at the onset of the channel-opening isomerization (Grosman et al., 2000). The existence of multiple, distinct C versus O structural pairs of binding sites suggests that there could be multiple, distinct isomerization pathways that culminate in multiple, distinct structural perturbations of the gate region. Accordingly, the C and O states in the schemes (Figures 1 and Figure 2 – figure supplement 1) represent conformational ensembles comprising several discrete (and stable) structures, {AC_LA_}⇄{AO_HA_}.

How much the elements of these efficiency-class ensembles might diverge regarding the structure and function at the pore domain is speculative. So far, there is no evidence in AChRs of agonist-dependent variations in channel conductance or ion selectivity. However, this possibility is worth mentioning because if even small difference are detected, such global conformational ensembles could provide insight into the mechanism of biased agonism in other receptors (Ehlert, 2018). That is, different outputs might reflect different efficiency classes.

Figure 4 shows the agonists grouped by efficiency. Ligands with cationic groups that occupy smaller volumes *in vacuo* appear in general to have greater efficiencies (Indurthi & Auerbach, 2021). For example, D increases in the series TEA-TMA-TriMA (0.40-0.54-0.60) and between BzTEA-BzTMA (0.29-0.51). In this regard it is perhaps relevant that the volume of the aromatic binding pocket appears to be smaller in LA versus HA configurations (Table 3 – source data 1). However, the relationship between agonist size, pocket volume and D is complex. For instance, Dec and SCh have the same-sized cationic center as ACh and CCh but lower (and different) efficiencies. The dynamic nature of loops, side chains, the agonist and water complicate such structure-activity relationships that could possibly be probed by using molecular dynamics simulations.

To fully understand agonist action, it will be necessary correlate changes in energy with those in structure in both catch (^A^C⇄AC_LA_) and hold (C_LA_⇄AC_HA_), the only rearrangements in activation that are agonist dependent (Figure 2 – figure supplement 1). Regarding catch, despite slow and agonist-dependent association into C, many of the agonists we examined diffuse freely into the active-O binding pocket with an association rate constant ∼4×10 M^-1^s^-1^, regardless of their potency (Nayak & Auerbach, 2017). This, and the fact that perfectly symmetric TMA binds to C ∼100 times more slowly than does asymmetric ACh, indicates that slow agonist association cannot be attributed a requirement for the ligand to first adopt a particular configuration. Rather, we conclude that making an AChR active constitutively (for instance, by adding mutations far from the neurotransmitter sites) enforces the catch rearrangement.

Indeed, the observation of diffusion-limited agonist association to O is explained if all of the component microscopic protein rearrangements that constitute gating are linked as an multi-stage LFER (Gupta et al., 2017). Accordingly, this extended LFER chain would demand that a spontaneous local rearrangement of any one intermediate stage will propagate in both directions, towards O and towards C. For example, a mutation at the extracellulartransmembrane interface might not only increase P_O_ constitutively (open the gate and allow ions to flow), but (because of microscopic reversibility) also cross the gating-binding divide and enforce catch and allow agonists to diffuse into the binding sites. The experimental observation of diffusion-limited binding to O (Grosman & Auerbach, 2000; Nayak & Auerbach, 2017) leads us to suggest that all of the microscopic domain rearrangements within the multi-stage bindinggating conformational cascade are linked energetically, as a single, extended LFER chain (Figure 2 – figure supplement 1).

In several other receptors, agonist association into resting bind sites is slower than diffusion (Jadey & Auerbach, 2012; Nayak & Auerbach, 2017). Why would natural selection generate a barrier for agonist entry into the C binding site? Our hypothesis is that the catch rearrangement is involved in molecular recognition. The forward catch rate constant (the agonist-dependent component of k_on_ to C) is correlated with agonist potency (Jadey & Auerbach, 2012; Jadey et al., 2011; Nayak & Auerbach, 2017). Because of the barrier, weak agonists that are present at the synapse (such as choline) and might interfere with physiological signaling at mature or developing synapses will be excluded from entering the neurotransmitter pocket. Another possibility is that catch, unlike diffusion, can be fine-tuned to set both CRC parameters and the synaptic current profile (see Methods).

The suggestion that catch (^A^C⇄AC_LA_) serves to select specific ligands for entry into the binding pocket places focus on the location(s) of the external agonist binding site(s) in ^A^C. In this regard it is worth noting that extracellular cations compete with agonists to slow k_on_ (Cs^+^>K^+^>Na^+^>Li^+^) (Akk & Auerbach, 1996), and that the mutation DE184Q eliminates this competition (Akk et al., 1999). However, none of the three states that comprise the catch-andhold LFER (^A^C⇄AC_LA_⇄AC_HA_) have been identified in structures.

Our understanding of the above results is incomplete. Regarding agonists, we do not know the structural basis for different efficiency classes, for instance ACh>Ebt>Var. Regarding mutations, we do not know the reasons D is reduced i) by DK145A for ACh but not Ebt, ii) by DD200A for all agonists except TMA, iii) by DG153S for all agonists and iv) by DY190A but not F for ACh. Regarding binding sites, we hope to learn the reason some agonists have greater efficiency at AChR fetal versus adult sites (Nayak et al., 2019), as well as agonist efficiency values at other nicotinic and non-nicotinic receptor neurotransmitter sites. We hope that a combination of structures, energy measurements and computational analyses will illuminate the molecular forces that underpin agonist efficiency and lead to a deeper understanding of how ligands promote conformational change in proteins.

## Materials and METHODS

<u>Expression.</u> Human embryonic kidney (HEK) 293 cells were maintained in Dulbecco’s minimal essential medium (DMEM) supplemented with 10% FBS and 1% penicillin–streptomycin (pH 7.4). Mutations were incorporated into AChR subunits using the Quickchange II site directed mutagenesis kit (Agilent Technologies, CA) according to manufacturer’s instructions. Sequence was verified by nucleotide sequencing (IDT DNA, I). AChRs were transiently expressed in HEK 293 cells by transfecting (CaPO_4_ precipitation) (Purohit et al., 2014) mouse D1 (GFP encoded between M3-M4)D D1D DD D subunits (3-5Dg total/ 35 mm culture dish) in a ratio of 2:1:1:1 for ∼16hrs. Most electrophysiological experiments were done 24-48 hrs post-transfection.

<u>Electrophysiology.</u> Single-channel currents were recorded in cell-attached patches (23° C). The bath solution was (in mM) 142 KCl, 5.4 NaCl, 1.8 CaCl_2_, 1.7 MgCl_2_, 10 HEPES/KOH (pH 7.4). High extracellular [K^+^] ensured that the membrane potential V_m_ was ∼0 mV. Patch pipettes were fabricated from borosilicate glass, coated with sylgard (Dow Corning, Midland, MI) to a resistance of ∼10 MΩ when filled with pipette solution (Dulbecco’s phosphate-buffered saline PBS) (in mM): 137 NaCl, 0.9 CaCl_2_, 2.7 KCl, 1.5 KH_2_PO_4_, 0.5 MgCl_2_, and 8.1 Na_2_HPO_4_ (pH /NaOH). Single channel currents were recorded using a PC505 amplifier (Warner instruments, Hamden, CT), low-pass filtered at 20 kHz and digitized at a sampling frequency of 50 kHz using a data acquisition board (SCB-68, National instruments, Austin, TX). For liganded activation experiments, agonists were added to the pipette solution at the desired concentrations. For unliganded activation experiments, we used pipettes and wires that were never exposed to agonists. To reduce the effect of channel block without affecting binding of agonist to the receptor, membrane potential (V_m_) was held at +70 mV when agonists were used (Jadey et al., 2011).

<u>Current analysis.</u> Analyses of the single-channel currents were performed by using QUB software (Nicolai, 2013). Single-channel currents occur in clusters when the opening rate constant is significantly large. For analysis, we selected clusters of shut/open intervals that appeared (by eye) to be homogeneous with regard to P_o_. We limited the analysis to intracluster interval durations and thus excluded sojourns arising from desensitized states (shut intervals between clusters >20 ms). The clusters were idealized into noise-free intervals after digitally filtering the data at 10-15 kHz (Qin, 2004). First, the idealized, intra-cluster intervals were fitted by a 2-state model, C⇄O. Then, additional nonconducting and conducting states were added, one at a time, connected only to the first O state, until the log likelihood failed to improve by 10 units(Qin et al., 1997). Cluster P_O_ at each agonist concentration was calculated from the time constants of the predominant components of the shut-(D_s_) and open-time distributions (D_o_): D_o_/(D_S_+D_o_). In this way, an equilibrium CRC was constructed as a plot of the absolute P_O_ (not normalized) versus the agonist concentration (see Figure 2).

<u>Equilibrium constants.</u> Two equilibrium dissociation constants comprise efficiency, K_dC_ and K_dO_ (Figure 1; Eq 3). These, and the fully-liganded gating constant L_2_, were estimated in two ways, with both methods producing the same results.

In the primary approach, K_dC_ and K_dO_ were estimated from CRC parameters (see Figure 2). The CRC was fitted by the Hill equation to estimate P_O_ (the high concentration asymptote) and EC_50_ (the agonist concentration that produces a half-maximum P_O_). Equilibrium constants were calculated from the reaction scheme pertaining to the main activation pathway (that assumes L_0_ and K_dO_ are negligible). Let x=[A]/K_dC_. For a one-site receptor (A+C⇄AC⇄AO) P_O_([A])= xL_1_/(1+x+xL_1_). For adult AChRs that have 2 equal and independent binding sites (A+C⇄ AC⇄A_2_C⇄A_2_O; red, Figure 1B), P_O_([A])= x^2^L_2_/(1+2x+x^2^+x^2^L_2_). Relating to CRC parameters, P_O_^max^ is the infinite-concentration asymptote, P_O_^min^ is the zero-concentration asymptote and EC_50_ is the concentration at which P_O_ is half P_O_^max^. Hence,

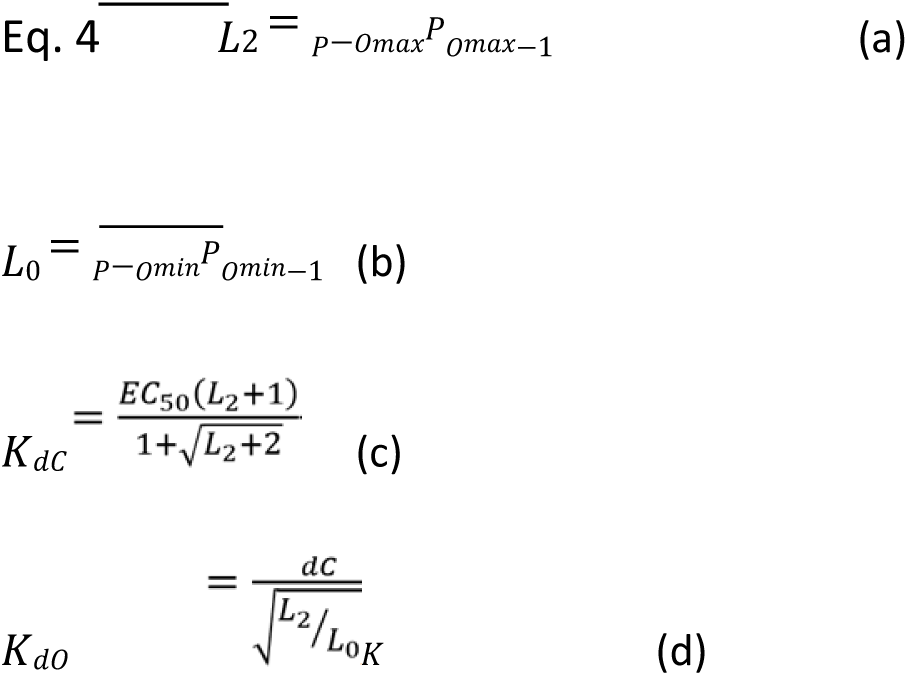

where K_dO_ is from Eq. 1. Also from Eq. 1 L_2_=L_0_^c^, so any change in L_0_ (see below) will change all 3 CRC parameters (P_O_^max^, P_O_^min^ and EC_50_) even if the equilibrium dissociation constant ratio K_dC_/K_dO_ 520 remains unchanged.

<u>Voltage, L_0_ and background mutations.</u> To reduce channel block by the agonists, the membrane was depolarized to +70 mV. To compensate for changes in D_S_and D_o_ caused by depolarization we added the background mutation εS450W. This residue is far from the binding site (M4 transmembrane region of the ε subunit), has no effect on K_dC_, and has equal but opposite effect on unliganded gating as does this extent of membrane depolarization (Jadey et al., 2011). L_0_ is x 10^-7^ at V_m_=-100 mV and reduced e-fold by a 60 mV depolarization (Nayak et al., 2012). Hence, we calculate that L_0_ is 5.2 x 10^-7^ at V_m_=-70 mV as well as in our experiments at V_m_=+70 mV plus εS450W.

In some conditions, for instance low efficacy agonists (Dec, TriMA, and BzTEA) and aD200A, the wt opening rate constant was small and single-channel clusters were poorly defined.

Accordingly, we added background mutations to facilitate P_O_ measurements (Tables 1 and 2). These were εL269F (located in the M2 helix) and εE181W (located in strand D9) that increase the L_0_ by 179- and 5.5-fold (1084-fold for the pair) without effecting K_dC_. First, we obtained the apparent L_2_ from the CRC P_O_ ^max^ (Eq. 4a). Second, we divided this value by the fold increase in L_0_ caused. By the background to obtain a corrected L_2_. Finally, agonist K_dC_ was estimated from EC_50_ and the corrected L_2_ (Eq. 4c).

<u>Catch.</u> The agonist association rate constant k_on_ to C (forward catch) determines the physiological response characteristics, as follows.

Experiments show that many agonists dissociate at about the same rate from C, k_off_∼15,000 s^-1^ (Jadey & Auerbach, 2012). The following 4 steps can be used a CRC from k_on_ if L_0_ and D are known *a priori*. Calculate i) K_dC_∼1.5×10^4^s^-1^/k_on_, ii) K_dO_ from K_dC_ and D (Eq. 2), and iii) L_2_ using L_0_ (Eq. 1), and iv) P_O_ and EC_50_ (Eq. 4). For example, using D=0.5, L_0_ =7.4×10 (at -100 mV) and k_on_=10^8^ M^-1^s^-1^ (for ACh) we calculatge K_dC_=150 DM, K_dO_=22 M, L_2_=33, P_O_ =0.97 and EC_50_=31 DM. With k_on_=5×10^6^ M^-1^s^-1^ (for choline), K_dC_ =3 mM, K_dO_=9 DM, L_2_=0.08, P_O_^max^=0.08 and EC_50_=6.8 mM. For both agonists, these all are a good approximation to be experimental values (Jadey et al., 2011). In AChRs, a measurement of k_on_ (forward catch), L_0_ and D are sufficient to estimate an agonist’s equilibrium properties. The procedure can be reversed, as k_on_ can be estimated from a CRC.

It is also possible to calculate the synaptic current time course from just k_on_. The channel-closing rate (b_2_) is approximately the same for many agonists, 2500 s^-1^ at -100 mV and 23 °C (Grosman et al., 2000). The synaptic current decay time constant (D) can be approximated as D=0.4(1+f_2_/2*k_off_). k_off_is 15,000 s^-1^ and f_2_ can be calculated from L_2_ (=f_2_/2500 s_-1_), calculated from k_on_, as above. Using the above k_on_ for ACh yields D=1.5 ms, close to the actual value. Fine tuning k_on_, the forward rate constant of the catch rearrangement (_A_C⇄AC; Figure 2) adjusts both the CRC and the synaptic decay.

<u>L_0_ for DD200A.</u> In wt adult AChRs, L_0_ is 5.2×10^-7^ at V_m_=+70 mV (Nayak et al., 2012) and is the same for all agonists. L_0_ has been reported previously for the mutations DK145A (Bruhova & Auerbach, 2017) and DG153S (Jadey et al., 2013). To estimate this L_0_ for DD200A, the pipette solution was free of any agonist and currents were measured at a membrane potential of −100 mV. The AChRs had added background mutations far from DD200 and each other (εL269F+εE181W+DV269A) that together increased unliganded activity substantially to allow cluster formation. Individually, these mutations increase L_0_ by 179- (Jha et al., 2009), 5.5- (Purohit et al., 2013) and 250-fold (Cymes et al., 2002) respectively (Figure 7 – figure supplement 1). Assuming no interaction (Gupta et al., 2017), the expected net increase in L_0_ for this background combination is the product, ∼2.5×10^5^. L_0_ was measured experimentally using this background plus DD200A, from the durations of intra-cluster intervals (see above). The unliganded opening (f_0_) and closing (b_0_) rate constants were estimated from the idealized interval durations by using a maximum-interval likelihood algorithm after imposing a dead time of 25 Ds and L_0_ was calculated from the ratio. Using a similar approach, L_0_ was estimated previously for each mutation shown in Figure 7.

<u>Statistical Information.</u> For single-channel CRCs, the midpoint and maximum (EC_50_ and P_O_) were estimated by fitting to monophasic hill equation (Po=P_O_ /(1+(EC_50_/[A]))) using GraphPad Prism 6 (GraphPad). A x-means cluster analysis algorithm (QUB online: qub.mandelics.com/online/xmeans.html) was used to define agonist (Figure 5) and mutant (Figure 7) groups, considering cluster as containing at least two data points. Optimal clustering was determined based on the Sum Square Residual (SSR) and corrected Akaike Information Criterion (AICc), for agonists (Figure 5A) SSR, 4.76 x 10^-3^ and AICc, -96.5 and mutations (Figure 7A) SSR, 2.41 x 10^-2^ and AICc, -396. Pearson’s correlation test was performed to determine correlation significance between the two variables logK_dC_ Vs logK_dO_ and logL_2_ Vs log[1/K_dC_] plots (Figures 5, 7). Since the aim is to measure the strength between the two variables and not the size of difference between the two groups, Pearson’s correlation was used over Cohen’s. The Pvalue (two-tail) and r^2^ value for agonist D/D classes (Figure 5) - 0.44/0.56, 0.019 and 0.96; 0.48/0.51, <0.0001 and 0.99; 0.57/0.41, <0.0001 and 0.99. Significance for clusters less than 3 data points (for D, 0.31 and 0.46) could not be determined. The P-value (two-tail) and r^2^ value for mutant D classes (Figure 7A) were 0.56, 0.0006 and 0.83; 0.51, <0.0001 and 0.98; 0.45, <0.0001 and 0.97; 0.41 <0.0001 and 0.93; 0.31, <0.0001 and 0.98.

<u>Agonists.</u> Abbreviations: acetylcholine (ACh), trimethyl ammonium (TRiMA), tetramethyl ammonium (Dwyer et al.), dimethylpyrrolidium (DMP), dimethylthiazolidinium (DMT), nornicotine (Kowal et al.), nicotinic (Nic), carbamylcholine (CCh), anabasine (Singh et al.), dimethylphenylpiperazinium (DMPP), benzyltrimethyl ammonium (BzTMA), choline (Cho), 3-hydroxypropyltrimethylammonium (3-OH), 4-hydroxybutyltrimethylammonium (4-OH), dimethylthiazolidinium (DMT), dimethylpyrrolidium (DMP), succinylcholine (SCh), decamethonium (Dec), epiboxidine (Ebx), epibatidine (Ebt), cytisine (Cyt), tetraethyl ammonium (TEA), tetramethyl phosphonium (TMP), decamethonium (Dec), varenicline (Var) and 599 benzyltriethyl ammonium (BzTEA). Agonists were from Sigma (St. Louis, MO) except DMP, DMT, 3-OH and 4-OH that were synthesized as described previously (Bruhova et al., 2013).

<u>Chemical potential.</u> The free energies DG_LA_ and DG_HA_ (Figure 1B), proportional to logK_dC_ and logK_dO_, are each a sum of a ligand-protein binding energy term and a chemical potential that incorporates the energy consequence of removing a ligand from solution. The magnitude of this term is indexed to a standard agonist concentration (M). In Eq. 2, we eliminated this potential from the accounting by first normalizing K_d_ values by those of a standard agonist,

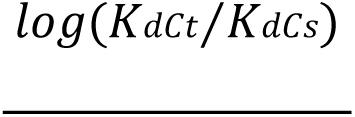

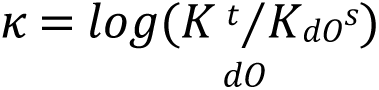

where the superscripts t and s denote the test and standard agonist. We used ACh, choline and Ebt as the standard agonist, with equivalent results.

Absent the chemical potential, D is a ratio of binding free energies, DG_catch_/DG_catch+hold_ (Figure 2B).

The position of AC_LA_ (D) in the catch-and-hold LFER (Figure 2) is analogous to the position of the transition state (D) in a rate-equilibrium free energy relationship of a single-step reaction (REFER, or Bronsted plot). In D (or Bronsted) analysis the intermediate state is an energy barrier, whereas in D analysis it is an energy well. D gives the fractional energy change at ‡, and D gives the fractional energy change at AC_LA_.

## SUPPLEMENTARY INFORMATION

**Figure 2 – figure supplement 1.**
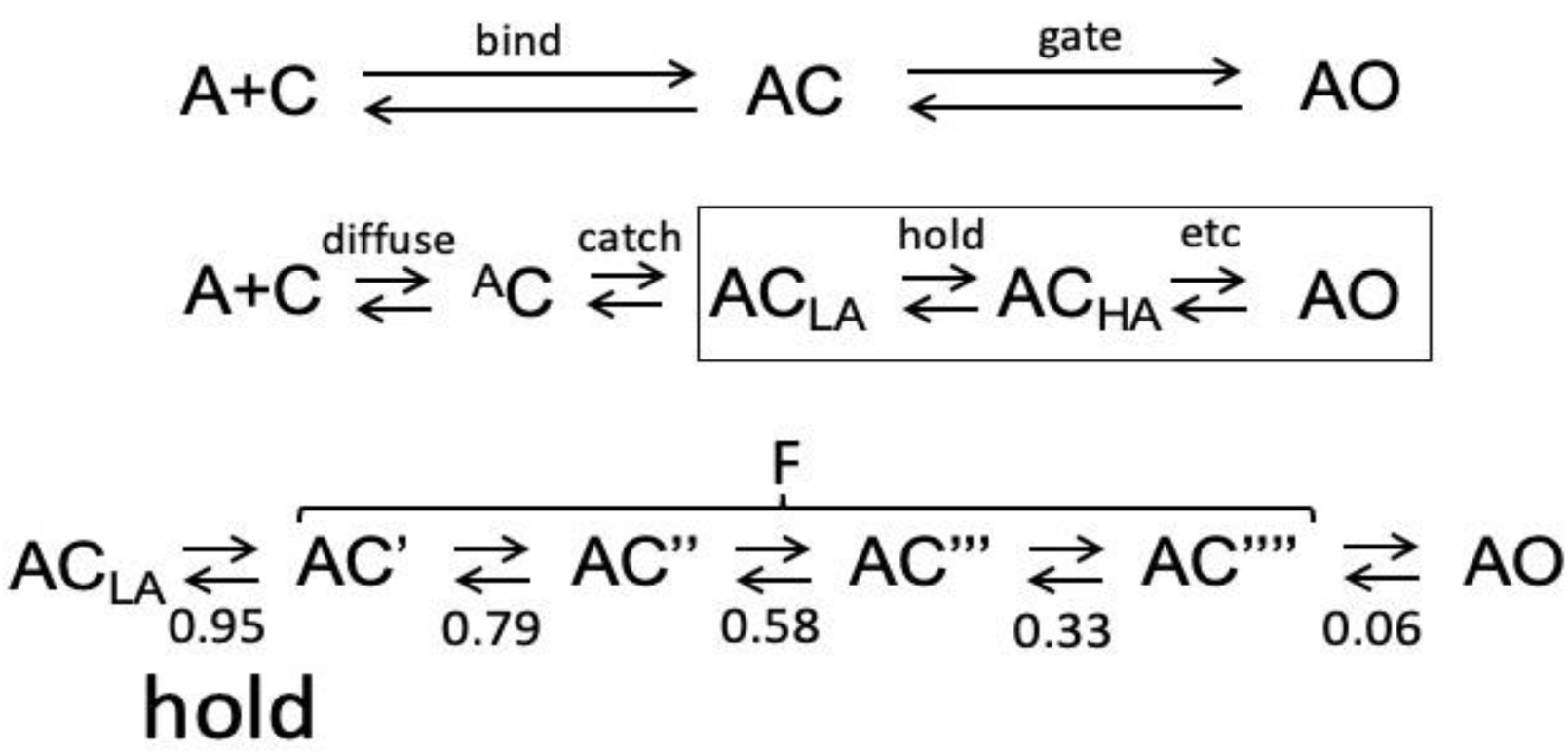
Intermediate states and transitions in AChR activation. C, closed channel; O, open channel; A, agonist (1-site receptor; see Fig. 1B). **Top**. *bind* and *gate* transitions bracket low affinity (LA) closed state AC (Delcastillo & Katz, 1957; Nayak & Auerbach, 2017). **Middle, l ro r.** In the bind step, the agonist reaches the neurotransmitter site (*diffuse*) and a local rearrangement generates the LA complex (*catch*), bracketing a state with very low affinity, ^A^C. In the gate step, another local rearrangement increases affinity (*hold*) and the rest of the protein isomerizes to O (*etc*), bracketing HA state ACHA (Jadey & Auerbach, 2012; Purohit et al., 2014). **Bottom.** Expansion of boxed region, middle. ACLA (left) is the only LA state; 4 short-lived (∼100 ns) HA states together constitute state F that is apparent in patch clamp recordings (*flip*; (Lape et al., 2008; Mukhtasimova et al., 2009)). Φ value of each microscopic transition is given below each set of arrows. The decreasing trend and longitudinal location in the protein suggests AChR gating is a conformational cascade that propagates from (l ro r) the agonist, extracellular domain, transmembrane domain, gate region and solvent (water+membrane) (Grosman et al., 2000; Purohit et al., 2013). Agonist free energy changes in the pre-catch and post-hold intermediate transitions are approximately the same for all tested ligands, so *catch* and *hold* are the only ones germane to efficiency (ΔGLA and ΔGHA; Eq. 2). Experimental evidence: *catch*, the association rate constant to C is i) slower than the limit set by diffusion, ii) correlated with agonist potency, iii) slower than to O (Nayak & Auerbach, 2017) and iv) can be highly temperature dependent (Gupta & Auerbach, 2011); *hold*, is high Φ values of agonists and binding site residues indicate a transition at the start of the gating sequence.

**Figure 3 – figure supplement 1.**
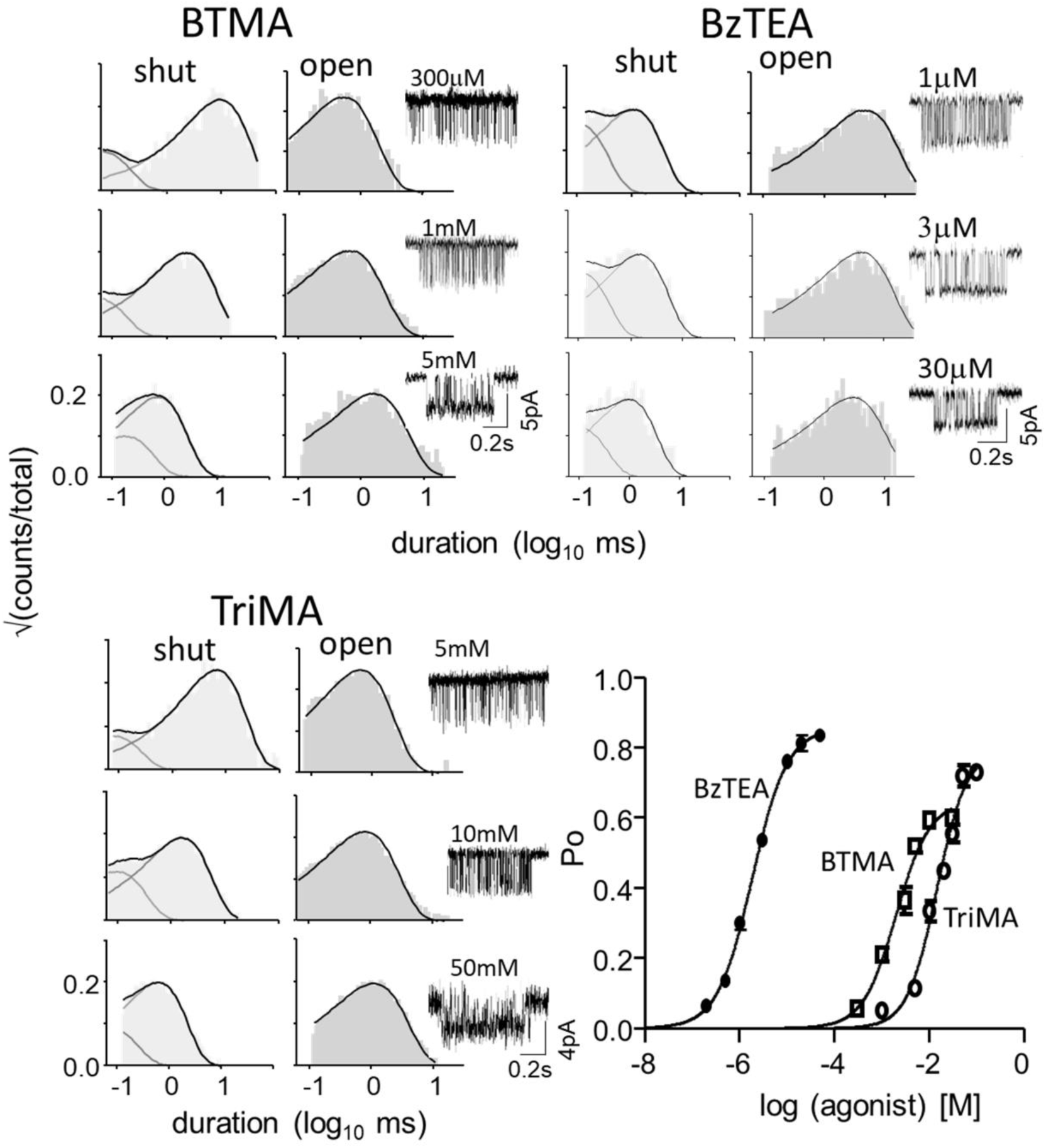
CRCs. Left, example histograms and clusters (see also Fig 3 and Fig. 4) Vm=+70, open is down. Efficiencies and background mutations are in Table 1. Symbols are mean±sem.

**Figure 6 – Source Data 1.**
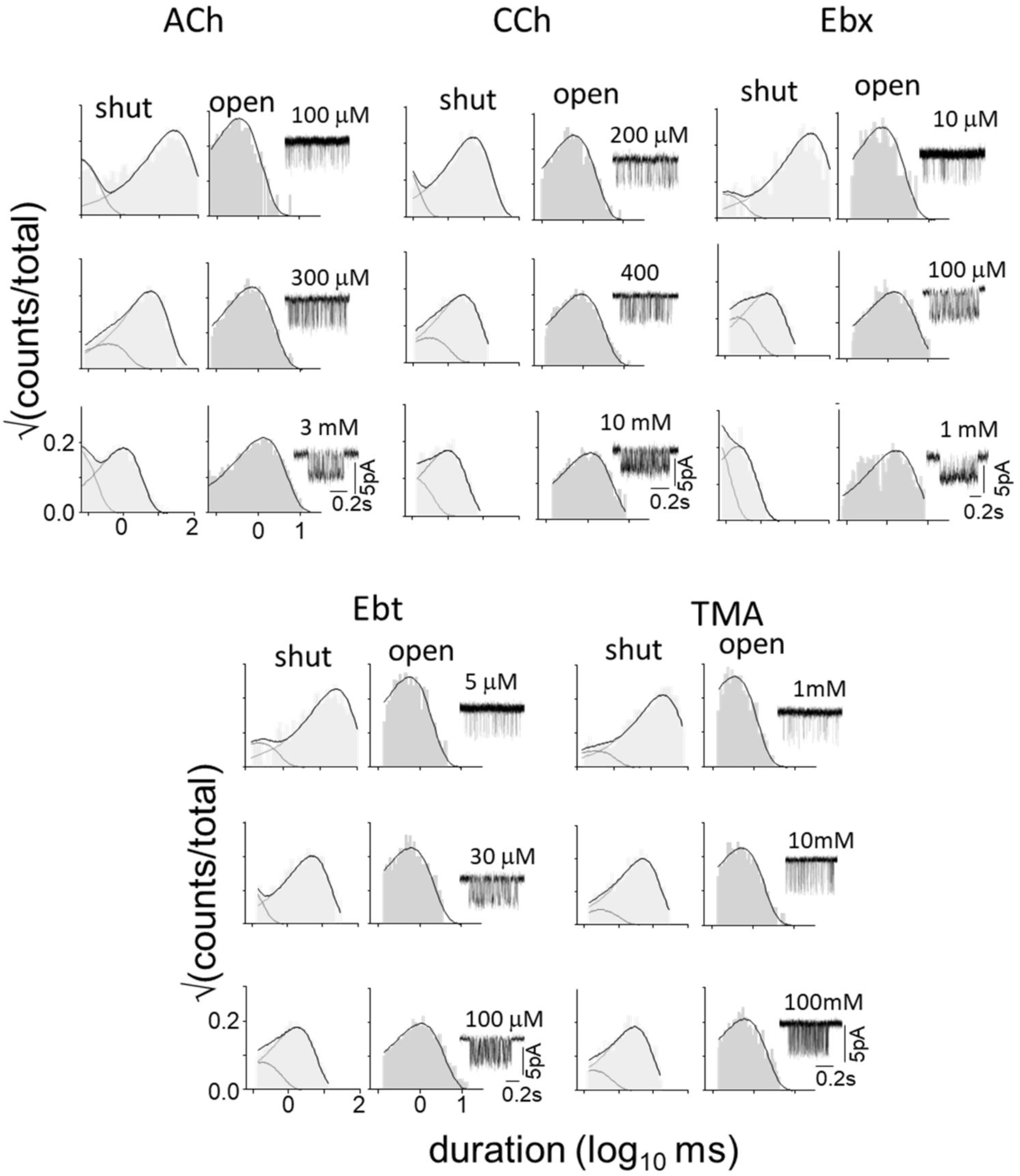
αK145A. Example histograms and clusters (agonist structures in Fig. 4). Vm=+70, open is down. CRCs in Fig. 6, efficiencies and backgrounds in Table 2.

**Figure 6 – Source Data 2.**
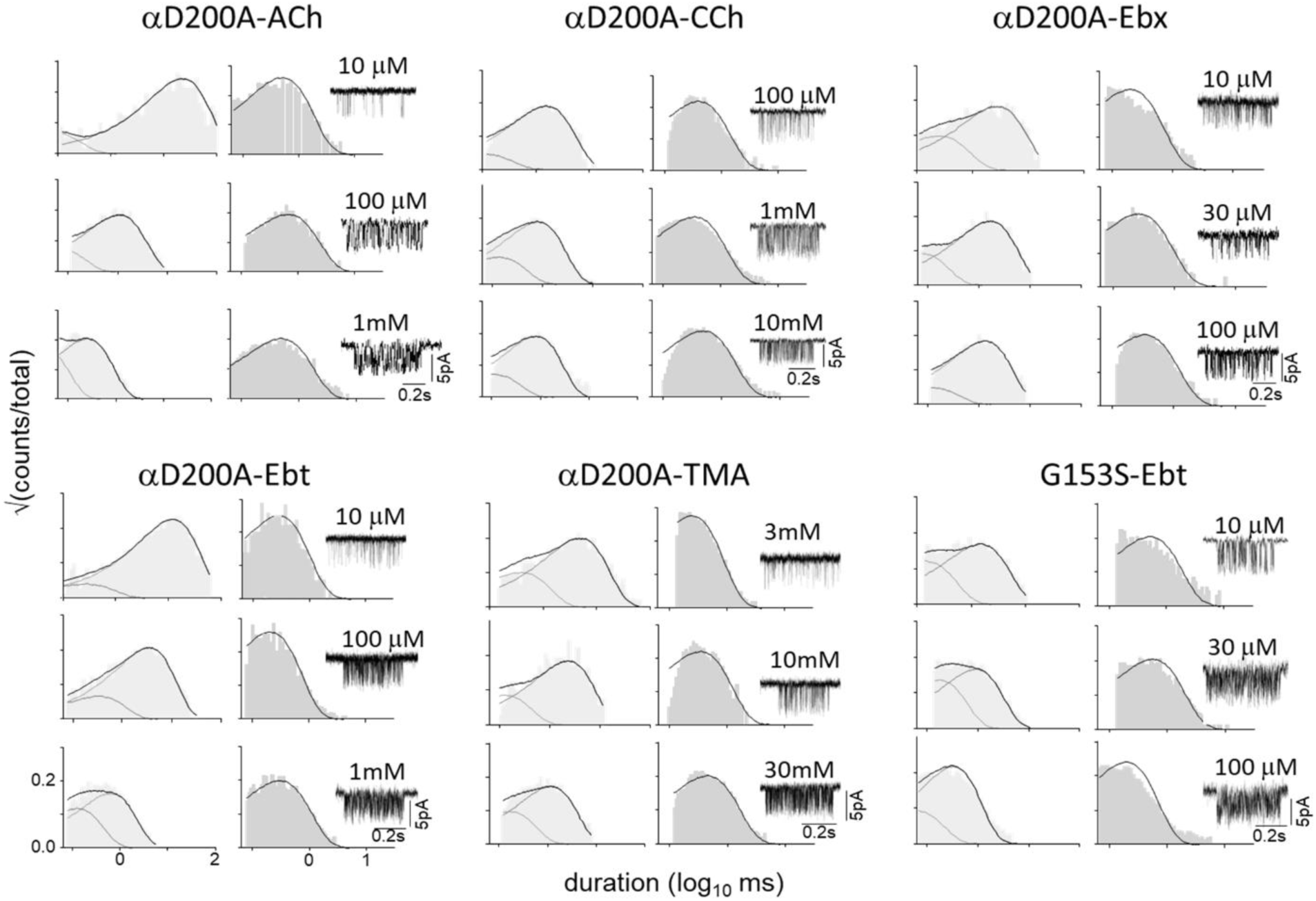
αD200A and αG153S. Example histograms and clusters (agonist structures in Fig. 4). Vm=+70, open is down. CRCs in Fig. 6, efficiencies and backgrounds in Table 2.

**Figure 7 – figure supplement 1.**
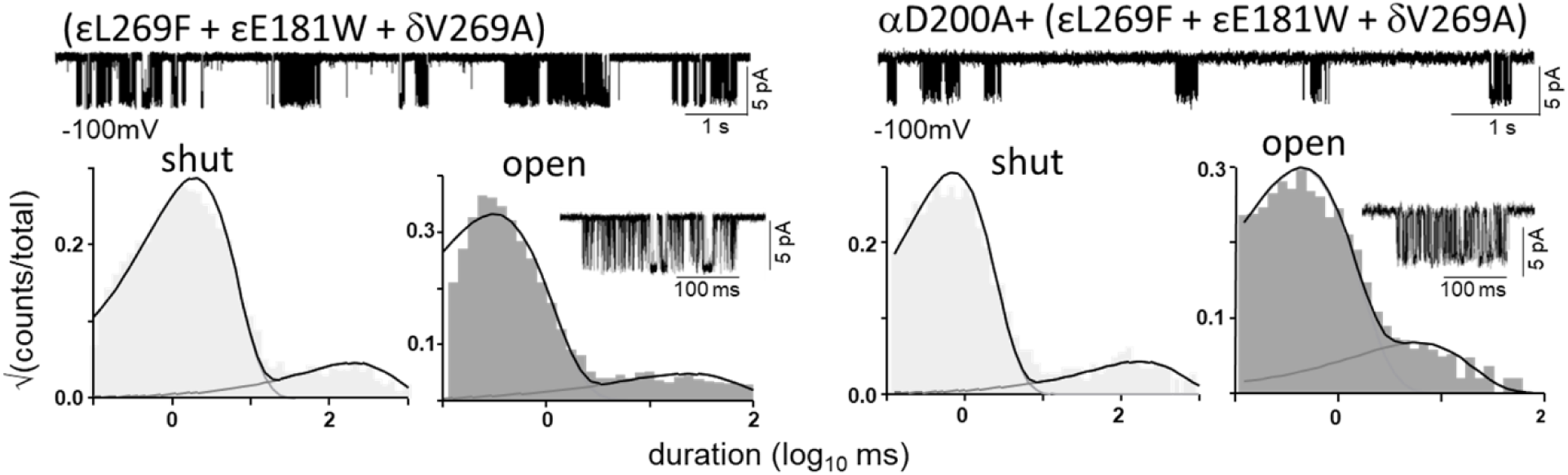
Estimating L0 (αD200). L, αD200 and R, αD200A. For both, the mutation combination εL269F+εE181W+δV269A was added to increase the WT L0 (7.4×10^-7^ at Vm=-100 mV) by a factor of 2.46×10^5^ (see Methods). Top, constitutive single-channel currents (no agonists; open in down). Bottom, example interval duration histograms and clusters (see Fig. 2). The observed L0= values were 0.17 (αD200) and 0.64 (αD200A). An A substitution at αD200 increases L0 by 3.76-fold. L0=5.2×10^-7^ in wt at Vm=-70 mV; (Nayak et al., 2012), so we estimate this membrane potential L0 for αD200A at 1.9×10^-6^. A similar measurement was made for every mutation in Tables 2 and S2.

**Table 1 – table supplement 1:** Agonist efficiency. KdC and KdO (Figure 2B) were calculated from CRCs (see Figure 3) after correcting L0 for the background (see Methods). c, coupling constant (KdC/KdO); η, efficiency (1-logKdC/logKdO). ^a^(Jadey & Auerbach, 2012), ^b^(Jadey et al., 2013), ^c^(Purohit & Grosman, 2006), ^d^(Indurthi & Auerbach, 2021).

| agonist | K<sub>dC</sub><br>(mM) | K<sub>dO</sub><br>(uM) | c | η |
| --- | --- | --- | --- | --- |
| ACh<sup>a</sup> | 0.17 | 0.03 | 6707 | 0.50 |
| Nor<sup>b</sup> | 0.20 | 0.04 | 4801 | 0.50 |
| CCh<sup>a</sup> | 0.54 | 0.17 | 3177 | 0.52 |
| Ana<sup>b</sup> | 0.23 | 0.04 | 6355 | 0.51 |
| Cho<sup>c</sup> | 4.10 | 15.10 | 297 | 0.50 |
| DMPP<sup>d</sup> | 0.48 | 0.13 | 3587 | 0.52 |
| 4OH<sup>d</sup> | 1.20 | 1.35 | 886 | 0.50 |
| 3OH<sup>d</sup> | 1.66 | 2.84 | 583 | 0.50 |
| Nic<sup>b</sup> | 1.00 | 0.92 | 1084 | 0.50 |
| TMA<sup>a</sup> | 1.48 | 0.62 | 2402 | 0.54 |
| DMT<sup>a</sup> | 2.73 | 2.76 | 989 | 0.54 |
| DMP<sup>a</sup> | 3.57 | 4.35 | 822 | 0.54 |
| Ebt<sup>d</sup> | 0.03 | 0.02 | 1698 | 0.41 |
| Cyt<sup>d</sup> | 0.07 | 0.10 | 650 | 0.40 |
| TEA<sup>d</sup> | 1.74 | 26.77 | 65 | 0.40 |
| TMP<sup>d</sup> | 0.36 | 1.47 | 244 | 0.41 |
| Var<sup>d</sup> | 0.06 | 0.33 | 171 | 0.34 |
| Ebx<sup>d</sup> | 0.11 | 0.05 | 2340 | 0.46 |

**Table 2 – table supplement 1:**
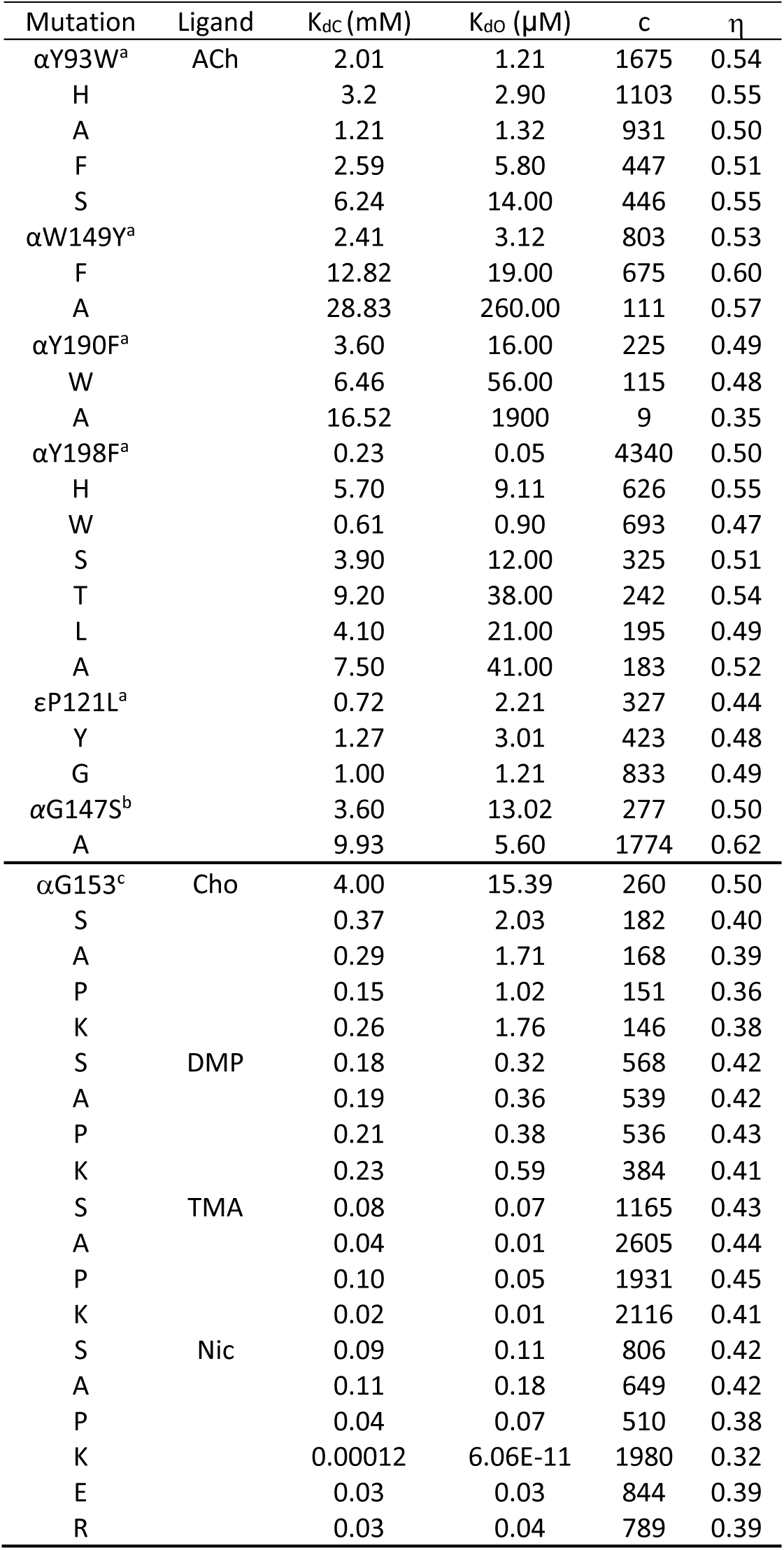
Mutant efficiencies. KdC and KdO were calculated from CRCs (see Figure 3); c, coupling constant (KdC/KdO); h, efficiency (1-logKdC/logKdO) (Eq. 3). ^a^(Purohit et al., 2014), ^b^(Purohit & Auerbach, 2011), ^c^(Jadey et al., 2013).

**Table 3 – source data 1.** Agonist binding-pocket volumes. Top row, subunit composition of the binding interface (see Figure 6 inset). Pocket volumes (A^3^) from homology models calculated as a pyramid with vertices at the centers of the five aromatic rings (Tripathy et al., 2019). LA, C and HA, O (Fig. 1). For all agonists and at all sites, the O pocket is smaller.

| Ligand | $\alpha-\gamma$ | | $\alpha-\delta$ | | $\alpha-\epsilon$ | |
| --- | --- | --- | --- | --- | --- | --- |
|  | LA | HA | LA | HA | LA | HA |
| ACh | 83.17 | 47.26 | 121.78 | 89.11 | 125.94 | 95.05 |
| CCh | 86.73 | 52.01 | 124.75 | 83.17 | 127.13 | 99.80 |
| TMA | 74.85 | 46.07 | 96.24 | 71.29 | 116.44 | 73.66 |
| Cho | 84.36 | 66.86 | 130.69 | 95.05 | 124.75 | 98.61 |

